# Identification of QTLs for Internode Length and Diameter Associated with Lodging Resistance in Rice

**DOI:** 10.1101/2025.10.21.683797

**Authors:** Qiuyun Lin, Ping Gan, Yujie Zhou, Yuehui Lin, Zhenyu Xie, Wei Hu

## Abstract

Lodging is a significant challenge in rice production, particularly for high-yielding cultivars. Previous research has shown that stem characteristics, such as internode length and diameter, are key determinants of lodging resistance. However, the genetic mechanisms controlling these traits remain poorly understood. To investigate the genetic basis of lodging-related stem traits, a recombinant inbred line (RIL) population was developed from a cross between Zhenshan 97 and C309, two rice varieties with contrasting stem architectures. A high-density genetic linkage map was constructed using 3,307 bin markers generated by next-generation sequencing. Comprehensive phenotypic analysis was conducted for heading date, internode length, and diameter. A total of 54 quantitative trait loci (QTLs) were identified across the genome. Importantly, 15 pleiotropic QTLs were detected, each affecting multiple internode-related traits. These pleiotropic loci include well-known functional genes such as *SCM2* and *Gn1a*, as well as several novel loci potentially involved in regulating stem strength and lodging resistance. Additionally, RNA-Seq analysis was performed to identify candidate genes associated with these traits. Among the 1,263 expressed genes within the 15 pleiotropic QTL regions, 183 differentially expressed genes were identified. *LOC_Os06g11130* was highlighted as a key gene for further investigation. Haplotype analysis and linkage disequilibrium analysis revealed significant haplotype variation in *LOC_Os06g11130*, further confirming its potential role in lodging resistance. This study enhances our understanding of the genetic architecture of internode traits associated with lodging resistance in rice. The identification of pleiotropic and major-effect QTLs provides valuable genetic resources for marker-assisted selection and genomic breeding, with the potential to improve lodging resistance in high-yielding rice cultivars. These findings contribute to the development of rice varieties that balance high yield potential with enhanced lodging resistance.

## 1 Introduction

Rice (*Oryza sativa* L.) is one of the most important staple crops globally, and its yield stability is essential for ensuring national food security. Lodging, caused by extreme weather conditions such as strong winds, heavy rain, or hail during the heading or grain-filling stages, is a major constraint on rice production (Pinthus, 1974; Shah et al., 2017a). It leads to the collapse of plants and panicles, damaging the vascular system and reducing grain filling, yield, and quality (Lang et al., 2012; Shah et al., 2019).

Environmental and agronomic factors, such as planting density, water and fertilizer management, and transplanting methods, significantly influence lodging resistance. For example, increased planting density results in reductions in internode diameter and wall thickness, which weakens lodging resistance (Shah et al., 2017b; Shah et al, 2019). Artificially-transplanted and mechanized planting rice exhibits stronger lodging resistance than direct-seeded rice (Terashima, 1997; Lei, 2014). Moreover, dry direct seeding improves grain yield and lodging resistance compared to flooded and wet direct seeding methods (Wang et al., 2021a).

Fertilization also plays a crucial role in lodging resistance. Proper nitrogen management enhances lodging resistance, while excessive nitrogen application can lead to increased plant height and a higher center of gravity, weakening stem strength and increasing lodging susceptibility (Corbin et al., 2016; Xiao et al., 2017; Pan et al., 2019).

The intrinsic determinants of lodging resistance are closely related to stem characteristics, including plant architecture, internode length, internode diameter, and wall thickness. These traits are regulated by quantitative trait loci (QTLs). In general, plant height is negatively correlated with lodging resistance (Yang et al., 1984; Niu et al., 2021). Additionally, the ideal plant architecture gene *IPA1*/*OsSPL14* results in reduced tillers, stronger culms, and improved lodging resistance (Jiao et al., 2010). Additionally, genes like *SCM2*/*APO1* and *SCM3*/*OsTB1* are involved in regulating panicle and stem strength, with variations in these genes contributing to different levels of lodging resistance (Takeda et al., 2003; Ookawa et al., 2010; Yano et al., 2015). Pyramiding *SCM3*/*OsTB1* with *SCM2* leads to increased stem diameter and wall thickness, resulting in stronger culms compared to single-gene lines. (Ookawa et al, 2010; Yano et al, 2015). *Gn1a* is a major-effect QTL that controls the number of grains per panicle in rice (Ashikari et al., 2005). The *gn1a* allele increases stem diameter while promoting crown root development, thereby enhancing the lodging resistance of rice (Tu et al., 2022). The recently mapped locus *qSCM4* has been shown to significantly increase wall thickness, thereby enhancing lodging resistance (Yang et al., 2023). A survey of the basal stem traits of 533 rice core germplasm lines revealed that differences in lodging resistance were primarily reflected in the internode diameter and wall thickness of the first and second internodes (Yuan et al., 2021). Recent studies have further revealed that culm diameter is a key determinant of lodging resistance. For instance, *STRONG1* (*MAP70*) was identified as a major locus controlling culm diameter through microtubule organization, where reduced *STRONG1* expression enhanced lodging resistance and yield by improving culm strength and panicle architecture (Zhao et al., 2025a). Similarly, *STRONG2* (*CSLA5*) was shown to regulate culm diameter and lodging resistance by promoting sclerenchyma development via cell wall biosynthesis, with natural allelic variation in its promoter region contributing to phenotypic difference (Zhao et al., 2025b).

Internode length and diameter in rice exhibit high heritability (Jeon et al., 2018; Zhao et al., 2021a; Akshay et al., 2024), making traits such as the length and diameter of the first, second, and third internodes important indicators of lodging resistance. Consequently, fine mapping of QTLs associated with internode-related traits is commonly used in genetic studies as an indirect method to identify QTLs governing lodging resistance (Zhu et al., 2008; Yadav et al., 2017; Jiang et al., 2019). To date, QTLs for internode length and diameter in rice have been predominantly mapped to chromosomes 1, 3, 4, 6, 8, and 11 (Kashiwagi et al., 2008; Kashiwagi, 2014; Yano et al, 2015; Zhao et al, 2021a; Zhao et al., 2023; Sowadan et al., 2024). Despite substantial progress in rice genetic research, studies specifically focused on lodging resistance remain relatively scarce.

In this study, recombinant inbred lines (RILs) were developed from parental lines with significant variations in internode length and diameter. The RIL population were subjected to next-generation sequencing to construct high-density genetic linkage maps. Several new QTL loci associated with stem traits were identified, including some with pleiotropic effects. The findings of this research will provide a foundation for the identification and functional validation of stem-related QTLs in rice, advancing our understanding of lodging resistance and offering valuable loci for molecular breeding applications.

## 2 Materials and Methods

### 2.1 Plant materials and population development

By crossing the indica rice varieties Zhenshan 97 (ZS97) and C309 to obtain F_1_ hybrids, one F_1_ plant was selected for self-pollination to develop the F_2_ generation, followed by single-seed descent to the F_8_ generation, resulting in a 209-line RIL population.

### 2.2 Trait measurement

To ensure the accurate identification of stable QTLs, field experiments were conducted at three farms in Danzhou City, Hainan Province, China, located in Nada Town, Wangwu Town, and Dongcheng Town, which were designated as three environments (R1, R2, and R3). In each environment, the same set of 209 lines of RIL and two parents was planted; however, their entry numbers were randomly rearranged across environments to minimize positional bias. For each line, ten plants were grown in a single row, with a row spacing of 16.5 cm and a plant spacing of 26.4 cm. Field management followed the same standard local practices for rice cultivation, including fertilization, irrigation, and pest control, in order to minimize non-experimental variation. For each family, heading date (HD) was recorded as the number of days from sowing to the emergence of the first panicle. To ensure uniform developmental stage, trait measurements were conducted 20 days after heading of RIL in April-May 2023. In all three environments (R1, R2, and R3), stem-related traits were measured by sampling three tillers per plant, and the mean values of three plants were used as the phenotypic value for each line. The internodes above the basal adventitious root were sequentially designated as the first, second, and third internodes. Internode length was recorded with a ruler, and internode diameter was measured twice at orthogonal positions using a vernier caliper, with the average value used as the phenotypic value. The correlation coefficients among these phenotypes were calculated and plotted using the corrplot package (Wei et al., 2017).

### 2.3 The DNA library construction, sequencing and genotyping

The fresh leaf samples of ZS97 and C309, and the RILs were sampled and submitted to the Novogene Company (https://en.novogene.com/) for Illumina HiSeq PE150 sequencing. The sequencing data generated was 1 G for the per RIL line and 5 G for each parent.

The raw data were filtered using the following criteria to obtain high-quality clean data: (1) reads with quality scores < 20e were discarded; (2) reads containing adapter sequences were removed; (3) reads with ≥ 10% unidentified nucleotides (Ns) were excluded; and (4) reads with > 50% of bases having a Phred quality score < 5 were eliminated. The clean reads were then mapped onto the rice genome of Nipponbare (Nipponbare, MSU version 7.0) (Kawahara et al., 2013) using BWA software (Li, 2013). The duplicate reads were removed by using SAMtools (version 1.7) (Li et al., 2009).

Population SNP calling was performed using GATK (GATK, version 4.1) (McKenna et al., 2010) (parameters: QD < 2.0, MQ < 40.0, FS > 60.0, SOR > 3.0, MQRankSum < −12.5, ReadPosRankSum < −8.0). Polymorphic sites were further identified using VCFtools (Danecek et al., 2011), with pre-filtering criteria applied to the parental lines: biallelic SNPs, no missing data, sequencing depth between 0.5× and 2× of the mean depth, and variants marked as “PASS” in the hard-filtering step. Subsequently, heterozygous genotypes in the RIL population were excluded to ensure accurate genotype calls for linkage map construction.

### 2.4 Linkage analysis and QTL mapping

A custom Perl script was utilized to eliminate redundant and distorted markers and convert the data into BIN format for further analysis (Wang et al., 2022). The genetic linkage map was constructed using the R/qtl package (Broman et al., 2003). QTL mapping was performed using the QTL.gCIMapping.GUI software (Zhang et al., 2020), with a genome scan step size of 1 cM and a significance threshold of LOD ≥ 2.5. Genetic distances were calculated using the adegenet package (Jombart, 2008), and a phylogenetic tree was constructed using ggtree (Yu et al., 2017). Part of the data visualization was performed using the ggplot2 package (Wickham, 2011).

### 2.5 Candidate gene screening within pleiotropic QTL regions

To identify candidate genes within pleiotropic QTL regions, the RNA-Seq expression data of Zhenshan 97 and Nipponbare stem tissues released by (Zhu et al., 2024) (https://biobigdata.nju.edu.cn/cart/download) were utilized. Differentially expressed genes within the regions were compared, and genes with |log2FC| ≥ 1 were selected as candidate genes. Additionally, gene annotation information was downloaded from the Rice Genome Annotation Project database (Kawahara et al, 2013). The annotation data were compared with published literature to further refine the selection and identify the most likely candidate genes.

### 2.6 Haplotype analysis

The SNP variation of the candidate gene *LOC_Os06g11130* of *PQ8* were obtained from the 3000 Rice Genome Project (Li et al., 2014). The haplotype analysis was conducted by geneHapR package (Zhang et al., 2023), and the linkage disequilibrium (LD) visualization was conducted by LDheatmap package (Shin et al., 2006). Phenotypic data for the haplotypes were obtained from the published data (Zhao et al, 2025b). The phenotypic differences between different haplotypes were assessed using *t*-tests to determine the statistical significance of the variation observed across haplotype groups.

## 3 Results

### 3.1 Phenotypic analysis of the RIL population

The HD, the length of the first, second and third internode (LFI, LSI and LTI), the diameter of the first, second and third internode (DFI, DSI and DTI) were measured and analyzed for two parental lines (ZS97 and C309) and their derived recombinant RIL population. ZS97 exhibits a shorter plant height relative to C309 (Figure 1A) and flowers earlier than C309 (Table 1). Upon removal of leaf sheaths and blades, five stem internodes were observed in ZS97, while six were present in C309 (Figure 1B).

**TABLE 1.** Descriptive statistics of HD and internode-related traits in RIL population across three environments (R1, R2, R3).

| Trait | Environments | Heritability (%) | CV (%) | ZS97 | C309 | RIL |  |  |  |
| --- | --- | --- | --- | --- | --- | --- | --- | --- | --- |
| | | | | | | Mean ± SD | Range | Skewness | Kurtosis |
| HD (day) | R1 | 91.6 | 9.1 | 69.0 ± 1.0 | 95.0 ± 1.0 | 77.8 ± 7.0 | 60.0–105.9 | 0.21 | 0.27 |
| | R2 | | 7.4 | 71.3 ± 1.2 | 93.7 ± 1.5 | 86.2 ± 6.4 | 68.1–100.8 | –0.00 | –0.55 |
| | R3 | | 8.1 | 70.0 ± 1.0 | 91.7 ± 1.5 | 79.3 ± 6.4 | 60.9–94.2 | 0.00 | –0.57 |
| LFI (cm) | R1 | 66.2 | 59.9 | 1.1 ± 0.1 | 0.7 ± 0.1 | 2.6 ± 1.5 | 0.1–12.5 | 1.60 | 5.50 |
| | R2 | | 53.7 | 1.0 ± 0.2 | 0.7 ± 0.1 | 2.6 ± 1.4 | 0.3–9.5 | 1.20 | 2.30 |
| | R3 | | 63.9 | 1.0 ± 0.4 | 0.7 ± 0.1 | 2.6 ± 1.6 | 0.1–12.7 | 1.90 | 7.56 |
| LSI (cm) | R1 | 65.6 | 38.9 | 4.1 ± 0.6 | 4.0 ± 0.2 | 6.2 ± 2.4 | 2.0–20 | 1.30 | 3.12 |
| | R2 | | 36.6 | 4.6 ± 0.4 | 4.1 ± 0.3 | 6.2 ± 2.3 | 1.7–15.8 | 0.90 | 1.04 |
| | R3 | | 39.6 | 4.2 ± 0.4 | 4.2 ± 0.4 | 6.2 ± 2.5 | 1.3–20 | 1.40 | 3.62 |
| LTI (cm) | R1 | 66.2 | 27.0 | 6.1 ± 0.1 | 9.5 ± 0.6 | 12.7 ± 3.4 | 4.9–30.0 | 1.00 | 3.17 |
| | R2 | | 27.4 | 6.2 ± 0.6 | 9.9 ± 0.3 | 12.6 ± 3.4 | 4.5–31.0 | 1.00 | 3.02 |
| | R3 | | 26.9 | 5.6 ± 0.2 | 9.4 ± 0.5 | 12.5 ± 3.4 | 4.5–31.0 | 1.00 | 3.56 |
| DFI (mm) | R1 | 57.2 | 12.1 | 4.69 ± 0.91 | 5.58 ± 0.60 | 6.43 ± 0.78 | 4.05–9.20 | 0.10 | –0.02 |
| | R2 | | 11.4 | 4.73 ± 0.56 | 5.59 ± 0.66 | 6.41 ± 0.73 | 4.00–8.50 | –0.10 | 0.00 |
| | R3 | | 12.0 | 5.01 ± 0.77 | 5.62 ± 0.10 | 6.4 ± 0.77 | 4.00–8.60 | –0.10 | –0.18 |
| DSI (mm) | R1 | 53.4 | 12.2 | 4.57 ± 0.82 | 5.24 ± 0.41 | 6.16 ± 0.75 | 3.70–8.50 | –0.10 | 0.05 |
| | R2 | | 12.0 | 4.53 ± 0.57 | 5.47 ± 0.47 | 6.15 ± 0.74 | 4.10–8.50 | 0.00 | –0.12 |
| | R3 | | 11.4 | 4.66 ± 0.66 | 5.48 ± 0.28 | 6.14 ± 0.70 | 4.35–8.20 | 0.00 | –0.21 |
| DTI (mm) | R1 | 52.2 | 13.2 | 4.30 ± 0.70 | 4.55 ± 0.19 | 5.52 ± 0.73 | 3.15–7.45 | 0.10 | –0.07 |
| | R2 | | 13.4 | 4.34 ± 0.75 | 4.60 ± 0.31 | 5.52 ± 0.74 | 3.55–8.00 | 0.30 | 0.02 |
| | R3 | | 13.9 | 4.24 ± 0.25 | 4.52 ± 0.23 | 5.53 ± 0.77 | 3.15–7.9 | 0.00 | –0.06 |

Note: HD, heading date. LFI, length of the first internode. LSI, length of the second internode. LTI, length of the third internode. DFI, diameter of the first internode. DSI, diameter of the second internode. DTI, diameter of the third internode. CV, coefficient of variation. R1, R2, and R3 represent the three field environments (Nada, Wangwu, and Dongcheng, respectively).

**Figure 1.**
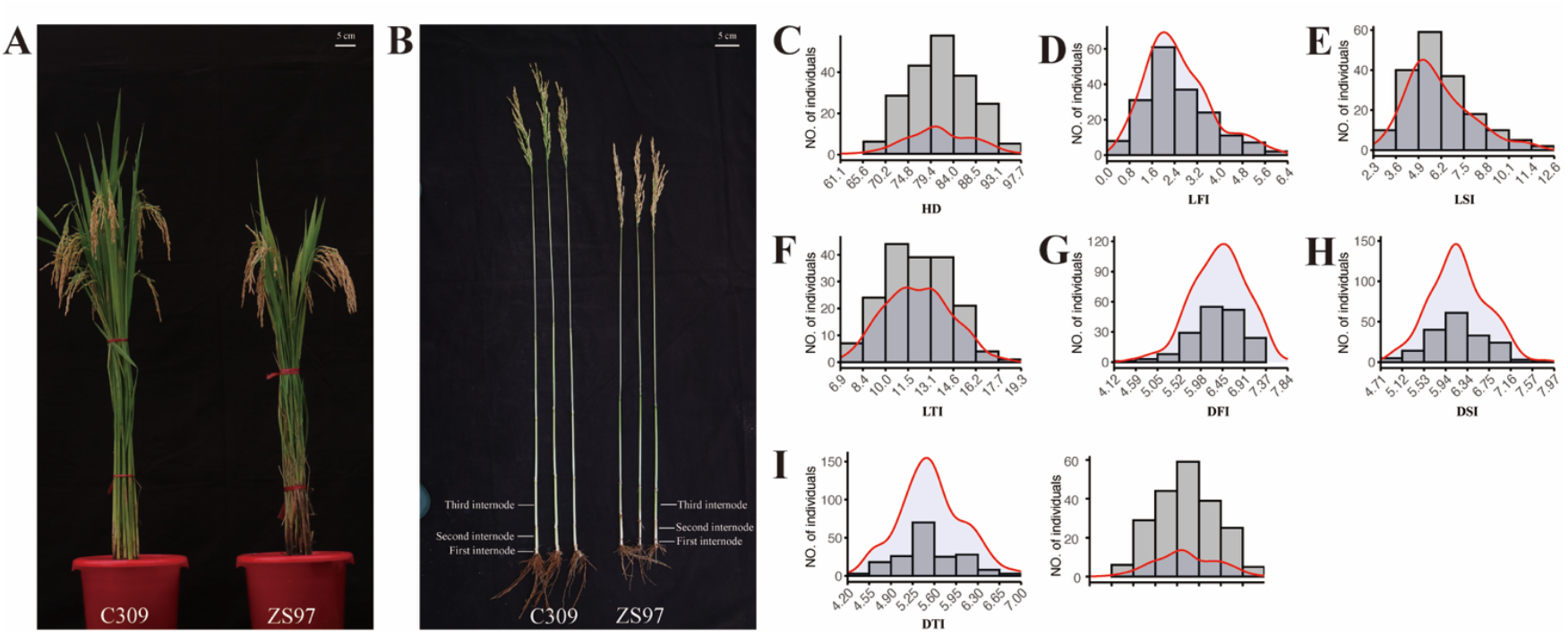
Phenotypic characterization and statistical analysis of the parental lines and RILs. (**A)**, plant architecture. (**B)**, Internodes of the parents. (**C**–**I)**, Histogram and kernel density estimation plots of phenotypic traits. All phenotypic values are presented as the mean across three environments (R1, R2, and R3).

The first internode of ZS97 was 0.3 cm longer than that of C309, while the second internode showed comparable lengths between the two varieties. However, the third internode of ZS97 was shorter than that of C309. In terms of internodes diameter across all three internodes, ZS97 exhibited smaller diameters compared to C309 (Figure 1B, Table 1). In both ZS97 and C309, the internode length increases progressively from top to bottom, whereas the internode diameter gradually decreases. Besides, the phenotypic values of the parents mostly fall within the variation range of the RIL population (Table 1). Between ZS97 and C309, all traits except for LSI showed significant differences based on Student’s *t*-tests (*P* < 0.05), whereas LSI did not differ significantly between the two parental lines.

The broad-sense heritability (H2) of HD and internode-related traits (LFI, LSI, LTI, DFI, DSI, and DTI) were calculated based on three repeats within each of the three environments (R1, R2, and R3). The heritability of HD was consistently high across all three environments, exceeding 90.0%, indicating that genetic factors played a predominant role in the variation of HD. For internode-related traits, the heritability of internode lengths (LFI, LSI, and LTI) ranged from 57.2% to 66.2%, reflecting moderate levels of genetic control, while the heritability of internode diameters (DFI, DSI, and DTI) was comparatively lower, ranging from 52.2% to 57.2%. The coefficients of variation (CV) varied across traits. HD exhibited relatively low CV values (7.4–9.1%), indicating stable expression. In contrast, LFI showed the highest CV (53.7–63.9%), reflecting large phenotypic variation. LSI and LTI had intermediate CV values (27.0–39.6%), while the CVs for internode diameters were smaller (11.4– 12.2%). These observations suggest that the RIL population captures the genetic diversity present in both parental lines and provides a broad spectrum of phenotypic variation for further QTL mapping to identify loci associated with internode length and diameter, which are important traits for lodging resistance in rice.

The correlation and significance analysis of internode-related traits reveals significant positive correlations between LFI, LSI, and LTI, as different internode lengths contribute to the overall structure of the plant. This correlation gradually weakens as the internode height increases. Notably, LFI is significantly correlated with DFI, but no correlation is observed between DSI and DTI. Internode lengths themselves are positively correlated with each other and significant. DFI is significantly positively correlated with all other phenotypic traits. Among these, the correlation coefficient between DTI and DSI is as high as 0.89 (Figure S1). These results suggest that DFI is crucial in influencing internode length and diameter.

### 3.2 Genotyping and genetic construction

A high-quality bin map of 3307 bins was constructed with 209 biparental RILs (Figure 2A, Table S1). The total length of the genetic map was 1,434.7 cM, with an average genetic distance of 0.44 cM per BIN. The distribution of BINs across chromosomes varied, with chromosome 1 containing the most BINs (374), and chromosome 12 containing the fewest (201). The chromosome lengths ranged from 86.3 cM on chromosome 12 to 168.6 cM on chromosome 1, with an average chromosome length of 119.6 cM. The average distance between adjacent BINs across chromosomes was 0.44 cM, with chromosome 8 showing the largest average distance at 0.51 cM, and chromosome 2 exhibiting the smallest average distance of 0.40 cM. The maximum gap observed between two adjacent BINs was 8.17 cM on chromosome 1, and the smallest gap was found on chromosome 10 at 1.76 cM (Table S1). The genetic linkage map of the RIL population was constructed based on the genetic distance (Figure S2B).

**Figure 2.**
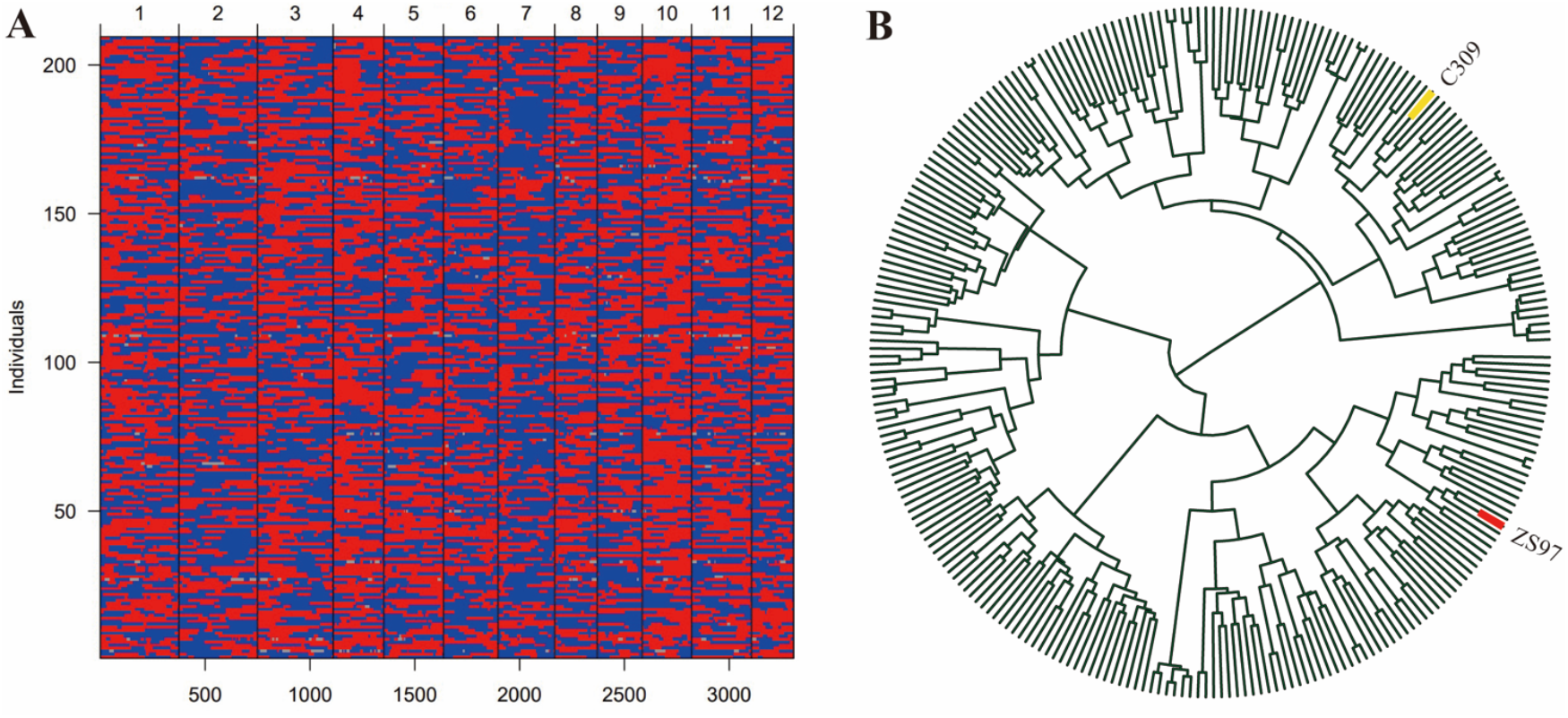
Bin map and phylogenetic tree analysis of the RIL population. (**A**), bin map, red plots represent allele of ZS97, while blue plots represent the allele of C309. (**B**), the phylogenetic tree of the RIL population.

In the phylogenetic tree constructed based on genetic distances, the RIL population and its parental lines clustered into distinct branches, with ZS97 and C309 located on two separate clades (Figure 2B). This indicates substantial genetic divergence between the two parents.

### 3.3 QTL for heading date

Given that heading date is closely associated with environmental adaptability in rice and is a highly heritable agronomic trait with relatively low environmental influence, QTL mapping was performed to identify potential loci related to heading date. This mapping also aims to utilize heading date as a marker to evaluate the accuracy of the genetic linkage map.

Through QTL mapping, four loci controlling HD were consistently detected across all three environments (R1, R2, and R3) (Table S2). These QTLs, designated *qHD1, qHD2, qHD3*, and *qHD4*, respectively (Figure 3). qHD1 was mapped to chromosome 3 at 3.4 cM, with a stable additive effect of approximately −2.6 and explaining 12.3–16.3% of the phenotypic variance (PVE). *qHD2* was located on chromosome 6 at ∼46–50 cM, with a positive additive effect (2.3–2.8) and explaining 12.0– 14.5% of the variance. *qHD3* was detected on chromosome 7 (∼56.7 cM) with smaller effects, accounting for 6.8–7.4% of the variance. *qHD4* was identified on chromosome 10 (79.5 cM), explaining 9.2–10.3% of the variance across environments. Among these loci, *qHD1* and *qHD2* showed the largest contributions, while *qHD3* and *qHD4* had relatively minor but consistent effects. Among them, *qHD1, qHD2*, and *qHD3* are likely associated with the cloned genes *OsMADS50, Hd1*, and *Ghd7*, which have all been previously reported to be related to heading date (Yano et al., 2000; Lee et al., 2004; Xue et al., 2008), with physical distances of 106.7 kb, 79.9 kb, and 950.7 kb, respectively. *qHD4* may represent a novel locus that has not yet been cloned.

**Figure 3.**
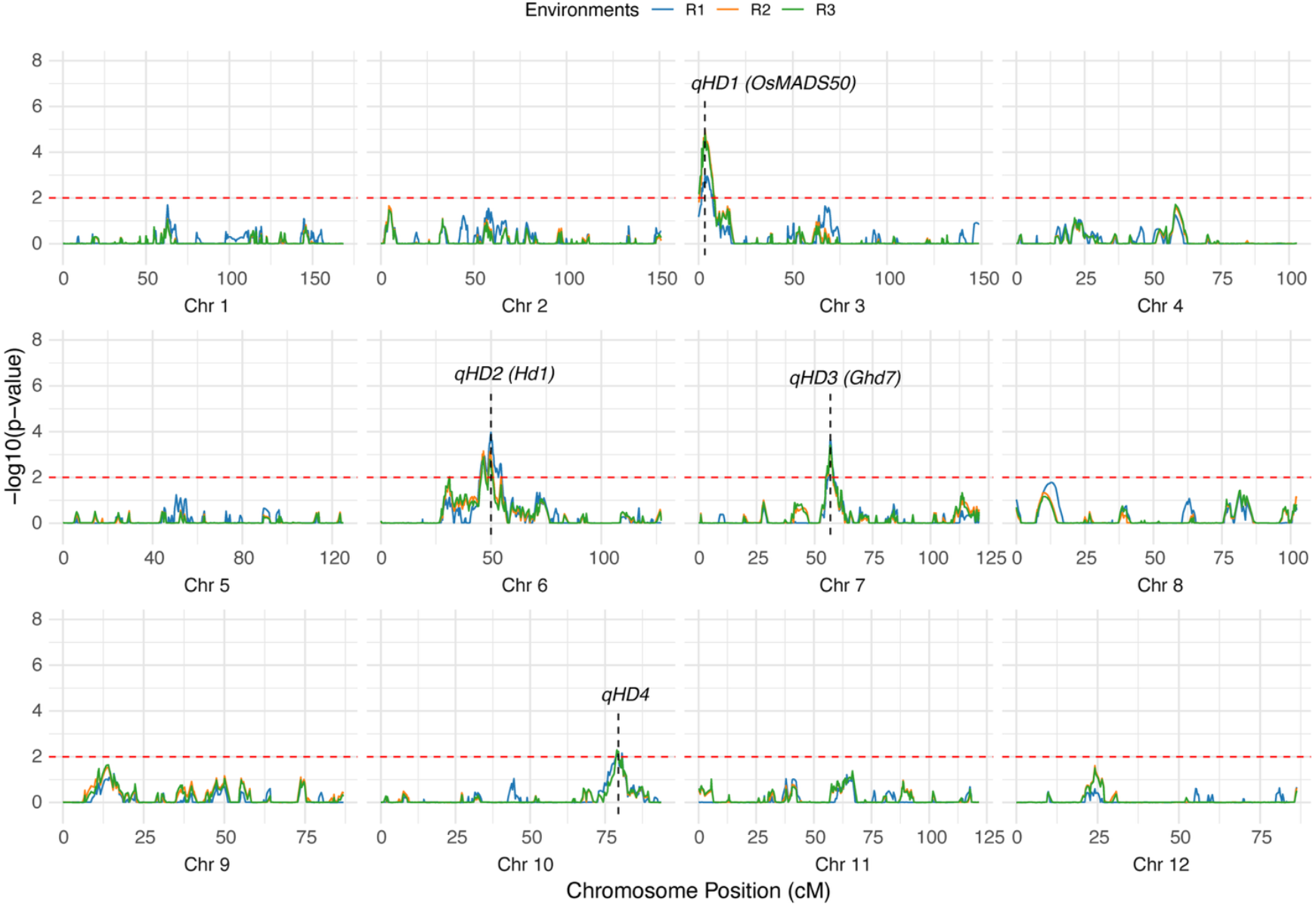
Mapping QTLs for HD by genome-wide CIM using the RILs. The dashed red line represents the threshold for significance (-log10(*p*-value) = 2). The labels indicate the positions of significant QTLs.

The repeated detection of these four QTLs across three environments highlights their stability and suggests that they represent reliable loci underlying heading date variation in rice. The consistent detection of these loci across multiple environments is likely associated with the high broad-sense heritability of HD (>90%), suggesting that the stable genetic basis of this trait enables reliable QTL identification. These results indicate that the genetic linkage map is robust and can be effectively used for dissecting stem-related traits.

### 3.4 QTL for internode-related traits

QTL mapping for internode-related traits identified multiple loci across the three environments (R1, R2, and R3), with results largely consistent with the high heritability of these traits and the observed phenotypic variation. For LFI, a total of seven QTLs were detected across different environments (Figure 4, Table S2). In R1, four QTLs (*qLFI_1, qLFI_2, qLFI_3*, and *qLFI_4*) were identified, with *qLFI_2* on chromosome 6 showing the largest effect (LOD = 6.6, PVE = 13.7%). In R2, two QTLs (*qLFI_5* and *qLFI_3*) were detected, both on chromosome 6, indicating partial overlap with the R1 results. In R3, five QTLs were identified, among which *qLFI_2* and *qLFI_4* overlapped with those in R1, while new loci such as *qLFI_6* (Chr4) and *qLFI_7* (Chr11) were also observed.

**Figure 4.**
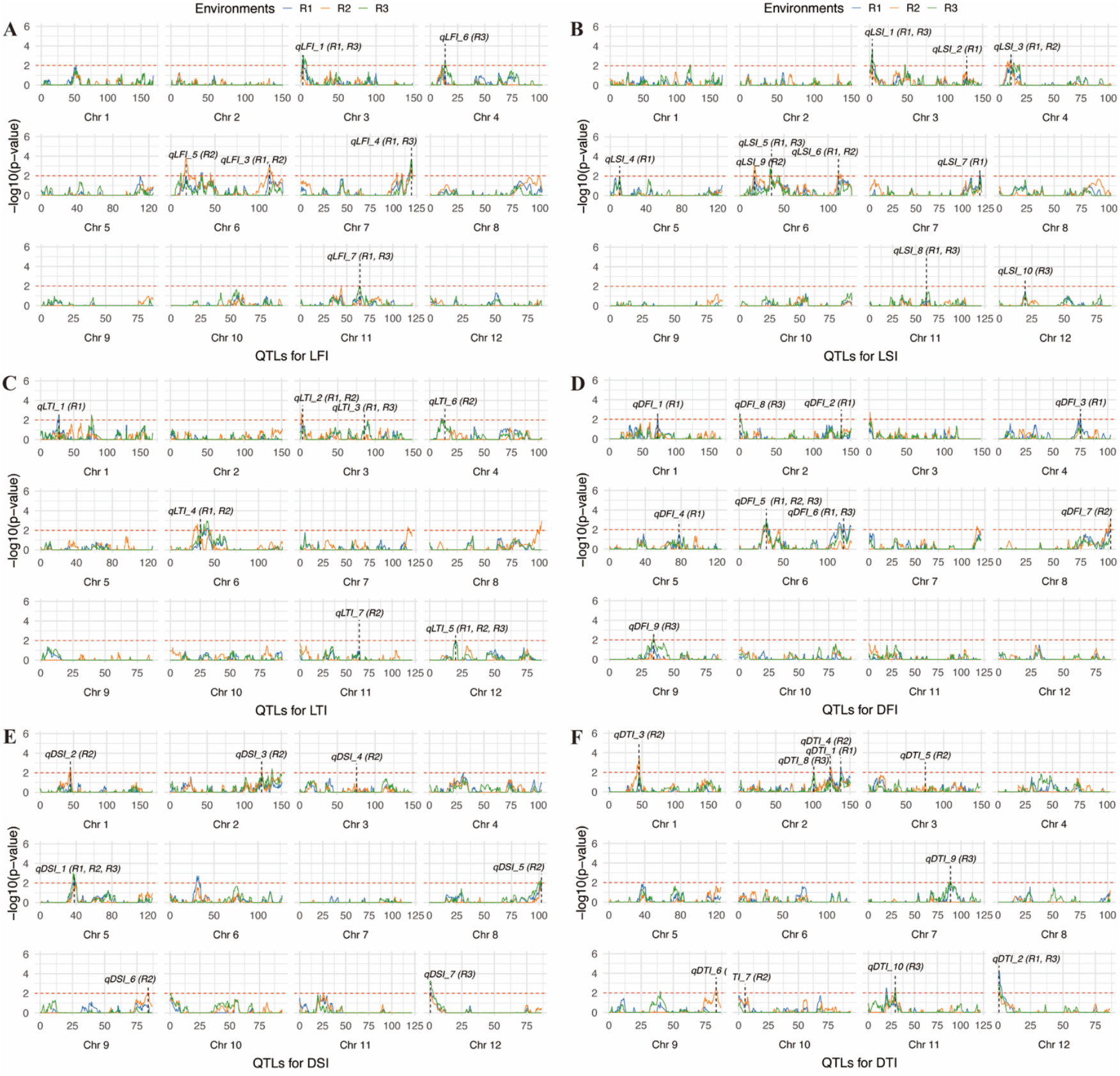
Mapping QTLs for LFI, LSI, LTI, DFI, DSI and DTI by genome-wide CIM using the RILs. The dashed red line represents the threshold for significance (-log10(*p*-value) = 2). The labels indicate the positions of significant QTLs. LFI, length of the first internode. LSI, length of the second internode. LTI, length of the third internode. DFI, diameter of the first internode. DSI, diameter of the second internode. DTI, diameter of the third internode.

For LSI, a total of ten QTLs were identified across the three environments. In R1, eight QTLs were detected, with *qLSI_5* (Chr6, LOD = 5.7, PVE = 7.9%) and *qLSI_6* (Chr6, LOD = 3.9, PVE = 4.6%) contributing the most. In R2, three QTLs were identified, including *qLSI_3* (Chr4), *qLSI_9* (Chr6), and *qLSI_6* (Chr6), of which *qLSI_6* overlapped with R1, confirming its stability. In R3, four QTLs were detected, including *qLSI_1* (Chr3), *qLSI_5* (Chr6), *qLSI_10* (Chr11), and *qLSI_8* (Chr12), with *qLSI_5* overlapping with R1, again indicating repeatability. Chromosome 6 harbors multiple QTLs for LSI detected across different environments, pointing to a key genomic region controlling this trait.

For LTI, seven QTLs were identified across three environments. In R1, five QTLs were detected, with *qLTI_4* (Chr6, PVE = 5.9%) being the most prominent. In R2, four QTLs were detected, including *qLTI_2* (Chr3) and *qLTI_5* (Chr12), which were also identified in R1, indicating stable detection. In R3, three QTLs were detected, with *qLTI_4* (Chr6) and *qLTI_5* (Chr12) again overlapping with previous environments. Thus, QTLs on chromosomes 3, 6, and 12 appear to play consistent roles in determining internode length across environments.

For DFI, nine QTLs were detected across environments. In R1, six QTLs were identified, among which *qDFI_5* (Chr6, LOD = 4.1, PVE = 7.7%) had the strongest effect. *qDFI_5* was also detected in R2 and R3, underscoring its stability. Additional loci were identified in R2 (*qDFI_7*, Chr8) and R3 (*qDFI_9*, Chr9), indicating environment-specific effects.

For DSI, seven QTLs were identified. R1 contained only one QTL (*qDSI_1*, Chr5), whereas R2 contained six QTLs spread across chromosomes 1, 2, 3, 5, 8, and 9, with *qDSI_1* (Chr5) overlapping with R1. In R3, two QTLs were detected, one of which (Chr5) overlapped with R1 and R2, confirming stability, while *qDSI_7* (Chr12) was specific to R3.

For DTI, ten QTLs were identified. In R1, two QTLs were detected, including *qDTI_2* (Chr12, PVE = 10.8%), which had the largest effect. In R2, six QTLs were identified, distributed across chromosomes 1, 2, 3, 9, and 10. In R3, four QTLs were detected, with *qDTI_2* (Chr12) reappearing and *qDTI_9* (Chr7) showing a strong effect (LOD = 3.7, PVE = 7.1%).

Overall, traits such as LFI, LSI, and LTI exhibited multiple QTLs with partial overlap across environments, consistent with their moderate heritability and relatively high phenotypic variation. In contrast, diameter-related traits (DFI, DSI, DTI), which exhibited lower coefficients of variation, showed fewer but more stable QTLs, such as *qDFI_5* on chromosome 6 and *qDSI_1* on chromosome 5. These results suggest that high-heritability traits with stable expression tend to yield reproducible QTLs across environments, while traits with higher environmental sensitivity tend to produce more environment-specific loci.

### 3.5 Pleiotropic QTLs

Pleiotropic QTLs, which influence multiple phenotypic traits (Hu et al., 2024), were identified in the current study. A total of 15 pleiotropic loci (*PQ1*–*PQ15*) were detected across the genome, several of which were concentrated in key genomic regions (Table S3, Figure 5). On chromosome 3, a strong pleiotropic interval (2.7–6.1 cM) was identified (*PQ4*), simultaneously regulating *qLFI_1, qLSI_1, qLTI_2*, and *qHD_1*, representing a hotspot where internode elongation traits overlap with heading date. Chromosome 6 emerged as a major hub of pleiotropy, with two distinct intervals: the first located at 30.7–35.8 cM (*PQ8*), which co-regulated *qDFI_5, qLFI_2, qLTI_4*, and *qLSI_5*, and the second spanning 111.8–118.5 cM (*PQ9*), which jointly controlled *qLFI_3, qLSI_6*, and *qDFI_6*. On chromosome 12, two pleiotropic loci were detected, including one at 0.7–1.2 cM (*PQ14*) that simultaneously regulated *qDSI_7* and *qDTI_2*, and another at 19.8 cM (*PQ15*) associated with *qLSI_8* and *qLTI_5*. Additional pleiotropic signals were also observed on chromosome 1 (*qDSI_2* and *qDTI_3* at 45.2 cM, *PQ1*), chromosome 2 (*qDSI_3* and *qDTI_4* at 123.4 cM, *PQ2*; *qDFI_2* and *qDTI_1* at 137.5 cM, *PQ3*),, chromosome 3 (*qDSI_4* and *qDTI_5* at 75.5 cM, *PQ5*), chromosome 4 (*qLFI_6, qLSI_3*, and *qLTI_6* at 9.4–13.8 cM, *PQ6*), chromosome 6 (*qLFI_6, qLSI_3* and *qLTI_67* at 16.7–17.2 cM, *PQ7*), chromosome 7 (*qLFI_4* and *qLSI_7* at 118.9–119.4 cM, *PQ10*), chromosome 8 (*qDFI_7* and *qDSI_5* at 102.3 cM, *PQ11*), chromosome 9 (*qDTI_6* and *qDSI_6* at 83.3–83.8 cM, *PQ12*), and chromosome 11 (*qLSI_10* and *qLFI_7* at 61.6–64.3 cM, *PQ13*). Collectively, these pleiotropic regions, particularly the strong clusters on chromosomes 3, 6, and 12, underscore the coordinated genetic regulation of stem elongation, diameter, and heading date. Importantly, the consistent detection of these QTLs across environments aligns with the high heritability estimates of these traits, suggesting that they represent stable and reliable hotspots. Such loci provide valuable targets for fine-mapping and molecular breeding aimed at enhancing lodging resistance and improving the adaptability of rice.

**Figure 5.**
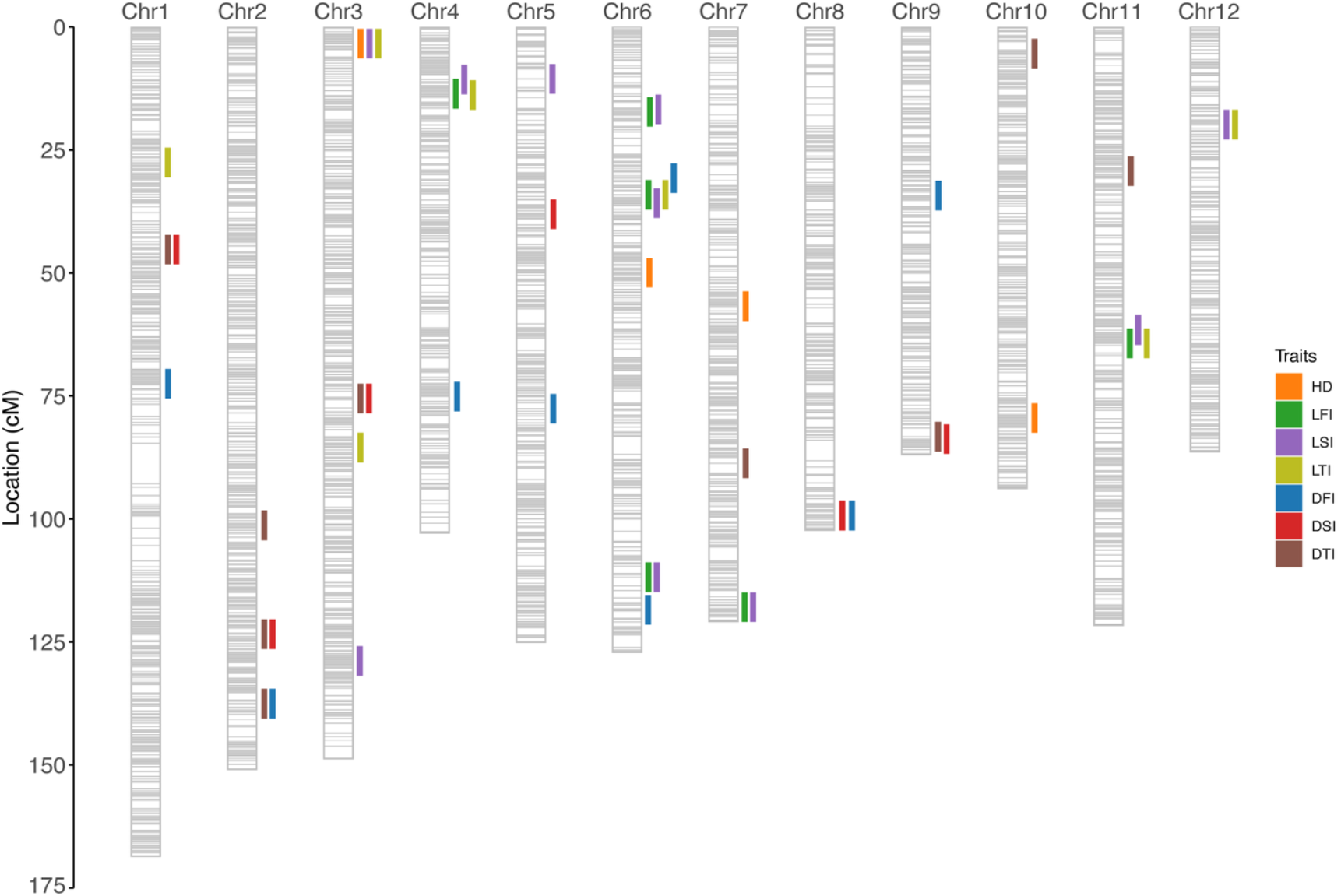
Genetic linkage map and QTLs location.

### 3.6 Candidate gene selection of pleiotropic QTLs

Screening candidate genes through the identification of differentially expressed genes is one of the commonly used approaches in QTL mapping and fine mapping, which offers the advantages of speed and convenience. Based on publicly available RNA-Seq data (Zhu et al, 2024), expression genes within pleiotropic QTL regions were selected. A total of 1,263 expressed genes were identified across 15 pleiotropic regions. Most of the pleiotropic QTL regions contained between 1 and 10 expressed genes, such as *PQ1, PQ2, PQ3, PQ5, PQ10, PQ11, PQ12, PQ14*, and *PQ15*. However, larger regions like *PQ4, PQ6, PQ8, PQ9*, and *PQ13* contained more than 20 expressed genes. Among them, *PQ6* contained the largest number of genes, with 436 genes. Differentially expressed genes were selected based on |log2FC| calculations, resulting in the identification of 183 differentially expressed genes across the two parents. No differentially expressed genes were found in *PQ1, PQ3, PQ5, PQ10, PQ11, PQ12*, and *PQ15*. This outcome may be attributed either to deviations in the defined QTL intervals or to the possibility that the functional differences of the genes within these regions are not driven by transcriptional variation. In other pleiotropic QTL regions, the number of differentially expressed genes was significantly lower compared to the total number of expressed genes. Specifically, *PQ2, PQ7*, and *PQ14* had fewer than five differentially expressed genes, while *PQ4, PQ6, PQ8, PQ9*, and *PQ13* contained 27, 64, 26, 16, and 43 differentially expressed genes, respectively (Table S3 and Table S4). The annotation information of the differentially expressed genes is provided in Table S4.

The selection of candidate genes was mainly accomplished based on their functional annotations and the roles of their associated families in rice stem development.

*LOC_Os03g03760* is annotated as a putative MYB family transcription factor. MYB transcription factors are well-established as upstream master regulators in the biosynthesis of secondary cell wall components (Zhao and Bartley, 2014), notably lignin and cellulose (Xiao et al., 2021). By controlling the synthesis of these structural polymers, this gene directly influences the mechanical properties, including its strength and wall thickness (Wang et al., 2014; Yang et al., 2014). Additionally, many MYB transcription factors have been reported to be associated with flowering time in rice (Wang et al., 2021b; Jin et al., 2025). Although *OsMADS50*, previously hypothesized as a candidate gene for *qHD_1*, is located within the same region, no differential expression was detected for *OsMADS50* in the expression analysis. Consequently, *LOC_Os03g03760* was selected as an important candidate for *PQ4* (*qLFI_1, qLSI_1, qLTI_2* and *qHD_1*), dissecting the genetic basis of internode length and diameter.

Finally, *LOC_Os04g12900* and *LOC_Os04g12980* are annotated as putative glucosyltransferases and UDP-glucosyl transferases. These enzymes are multifunctional, playing roles in the synthesis and modification of cell wall polysaccharides (cellulose and hemicellulose) essential for cell wall integrity (Zabotina et al., 2008; Wang et al., 2021c). Additionally, they may be involved in the glycosylation and inactivation of plant hormones (Škyvarová et al., 2025), thereby influencing cell division and elongation. Their dual roles in cell wall metabolism and hormone regulation position them as crucial candidate genes for elucidating stem structure and growth, and they are therefore considered potential candidates within *PQ6*.

The loci *LOC_Os06g11130* and *LOC_Os06g11135*, predicted to encode the gibberellin receptor GID1L2, were identified as differentially expressed genes within the *PQ8* region. This receptor is a central component of the gibberellin (GA) signaling pathway. GA signaling mediated by GID1 receptors is crucial for stem and internode elongation. In rice, downregulation of *GID1L2* family members in the *sped1-D* mutant causes shortened branches, while overexpression restores elongation, highlighting their role in culm development (Jiang et al., 2014). Similarly, mutations in *GID1L2* genes underlie dwarfism in watermelon (Liu et al., 2022), and evolutionary studies show that the GID1/DELLA module originated in vascular plants (Hirano et al., 2007), underscoring its conserved function in regulating stem growth and architecture across species. The role of GID1L2 in mediating the GA response makes it a primary candidate for understanding the genetic control of internode related-traits.

For other pleiotropic QTL regions, the differentially expressed genes have not yet been reported in plants, and therefore no candidate genes could be identified at this stage.

### 3.7 Genetic diversity and LD Analysis of *LOC_Os06g11130*

*PQ8* is the strongest QTL explaining phenotypic variation, controlling both internode length and diameter. Therefore, we focused on this region in the linkage disequilibrium and haplotype analysis. Based on its strong association with these traits, *LOC_Os06g11130* (AK070299) was selected for further analysis. This gene plays a significant role in the elongation of pedicels and secondary branches, as demonstrated by its reduced expression in the *sped1-D* mutant (Jiang et al, 2014). The overexpression of *LOC_Os06g11130* in *sped1-D* plants restored the normal panicle phenotype, indicating its involvement in regulating plant architecture. Given its critical role in gibberellin signaling and its potential impact on internode length and diameter, *LOC_Os06g11130* was prioritized for investigation in both LD and haplotype analysis to explore its genetic variation and association with agronomic traits.

SNP variations in *LOC_Os06g11130* were screened from the 3000 Rice Genome Project (Li et al, 2014), identifying a total of 6 SNP variations (located in 5835826, 5836263, 5836330, 5836463, 5836905, and 5836978) in the UTR and CDS regions. Combined with the phenotypic data from Zhao et al (2025b), 201 varieties that contained both SNP variations and phenotypic data were selected for further analysis (Table S5).

Haplotype analysis revealed a total of 7 haplotypes. The number of varieties for haplotypes 1-4 were 110, 47, 19, and 19, respectively (Figure S3A). The remaining three haplotypes, with fewer than five varieties (Table S5), were not displayed. For Hap1–4, there are only four SNP variation sites (5835826, 5836263, 5836463, and 5836905). SNP variation sites in 5836330 and 5836978 are only found in Hap6 and Hap5, respectively. Hap1 is predominantly found in indica (68.2%) and temperate japonica (29.1%), while hap2 and hap3 exhibit distinct subgroup characteristics. Hap2 is mainly associated with indica (97.9%), whereas Hap3 is exclusive to temperate japonica (100%). Hap4 is primarily found in indica (57.9%) and tropical japonica (36.8%) (Figure S3A). According to the haplotype analysis results (Table S5), the allele of *LOC_Os06g11130* in ZS97, recorded as CX133 in the 3000 Rice Genome Project database, is classified as Hap1. Although C309 is not included in the database, analysis of next-generation sequencing data from both parents indicates that the allele of *LOC_Os06g11130* in C309 is assigned to Hap2.

Significance testing between haplotypes revealed that Hap3 showed significant differences compared to all other haplotypes, while no significant differences were found between Hap1, Hap2, and Hap4, indicating that Hap3 may represent a distinct type. These results suggest that Hap3 is an inferior haplotype, which aligns with the findings of haplotype analysis (Figure S3B). A significant linkage disequilibrium (r^2^ > 0.8) was detected between the variation sites 5836330 and 5836463, suggesting that they are rarely separated by recombination events during the breeding process (Figure S3C).

## 4 Discussion

### 4.1 Environmental influences on lodging resistance

Lodging–the collapse of rice plants under forces like wind or rain–has long been recognized as a major constraint on rice production (Pinthus, 1974). The resistance to lodging is determined by a combination of genetic traits and environmental or agronomic factors (Zhang et al., 1999). Directly measuring lodging in field conditions is challenging because lodging occurrence is highly influenced by variable weather and field conditions. Pushing resistance of the stem is one direct indicator of lodging resistance (Kashiwagi et al, 2008; Torró et al., 2011; Kashiwagi, 2022), and a previous study identified the QTL *PRL4, prl5* and *OsPSLSq6* that improves pushing resistance of the lower stem (Kashiwagi and Ishimaru, 2004; Zhao et al., 2021b; Kashiwagi, 2022). In our preliminary measurements, the bending force required to bend the rice plants (pushing resistance) to 45 degrees at half of the plant height was measured in the RIL population as a proxy for lodging. However, no corresponding QTL was detected. The likely reason is the high phenotypic variation in this trait within the population, caused by differences in heading time, soil moisture, and maturity among lines. This difficulty highlights how environmental heterogeneity can obscure genetic effects on lodging when directly measuring it. Therefore, measuring indirect traits as indicators of lodging resistance in rice may be a more effective approach. This is because, according to current reports, the resistance to lodging in rice is highly correlated with traits the first to third internodes (Jiang et al, 2019; Zhao et al, 2021a).

In this study, we observed the impact of environmental factors on the stability of QTL detection for rice traits across three distinct environments (R1, R2, and R3). Some QTLs, such as the QTLs for heading date, *qDFI_5, qDSI_1*, and *qLTI_5* were consistently detected across all environments (Table S2), suggesting these loci are stable and have a robust genetic basis. Similarly, QTLs like *qLFI_2* and *qLTI_4* were identified across multiple environments, indicating their stable expression in controlling internode-related traits. These results demonstrate that certain genetic factors have relatively low environmental sensitivity, making them reliable targets for breeding programs.

Despite maintaining uniform planting spacing and consistent management of fertilizers, pesticides, and pest control across the three environments, environmental factors likely influenced the stability of QTL detection. Differences in plant height and heading date among the materials could have affected the growth of neighboring plants, leading to variations in phenotypic expression. For instance, disparities in developmental stages may have resulted in stronger genetic effects for some QTLs in certain environments, while these effects were weaker in others due to interplant competition. Additionally, differences in soil fertility between farms could have contributed to varying results in QTL detection. Factors such as soil moisture, nutrient content, and other physical and chemical properties may have influenced plant growth and, consequently, the expression of QTLs. Therefore, despite our efforts to standardize environmental conditions, these uncontrollable factors may have played a role in affecting the stability and genetic effects of the identified QTLs.

### 4.2 Genetic dissection of heading date

In this study, we explored the genetic basis of lodging-related internode traits using a recombinant inbred line (RIL) population derived from ZS97 and C309, two rice varieties with distinct plant architectures. ZS97 is shorter and more lodging-tolerant, while C309 is taller with thicker internodes. The RIL population exhibited broad phenotypic variation, with most trait values exceeding the parental ranges (Table 1), indicating segregation of diverse alleles. Using a high-density bin map with 3,307 markers, we detected several QTLs for heading date and internode-related traits (Figure 4, Table 2). This confirms sufficient genetic resolution for dissecting lodging-associated traits.

Heading date is a critical developmental trait that affects both the growth duration and plant size, which are essential factors in lodging resistance. In this study, we identified four QTLs for heading date across three environments (R1, R2, and R3) (Table S2). These QTLs, *qHD1, qHD2, qHD3*, and *qHD4*, were consistently detected across environments, underscoring their stability and genetic relevance. These QTLs are likely associated with genes previously reported to affect flowering time, including *OsMADS50, Hd1* and *Ghd7*, which play pivotal roles in regulating heading date and plant stature (Yano et al, 2000; Lee et al, 2004; Xue et al, 2008). Notably, variation in heading date can influence the duration of vegetative growth and the plant’s overall size, which directly affects its lodging resistance.

Breeding for lodging resistance requires balancing the extension of the vegetative growth period to enhance yield potential while simultaneously strengthening the stem to avoid lodging. Recent studies suggest that optimizing pre-heading growth can improve lodging resistance without compromising yield, as manipulating heading date allows breeders to adjust the timing of stem development to strengthen culm sturdiness (Guo et al., 2021). In our study, the identification of pleiotropic genes influencing both heading date and stem strength emphasizes the importance of developmental timing in lodging resistance. *qHD3* and *qHD4*, which are located on chromosome 7 and chromosome 10, respectively, provide new targets for breeding and gene editing to improve lodging resistance while maintaining high yield potential. *qHD4*, a novel QTL, accounted for approximately 10.3% of heading date variation and may represent a previously unidentified locus related to heading time.

### 4.3 Pleiotropic loci underlying multiple lodging-related traits

Our mapping revealed numerous QTL for individual internode lengths. Several loci were repeatedly detected across multiple internodes, indicating common genetic determinants for internode elongation. On chromosome 6, for example, a region around 30.7–35.8 cM harbored the largest-effect QTL for LFI, LSI, LTI and DFI (*PQ8*: *qDFI_5, qLFI_2, qLTI_4* and *qLSI_5*), each with LOD scores >3 and individually explaining about 5.9%–13.7% of the variation. Another notable region was on chromosome 7 at 118–119 cM, which affected both first and second internode length (*PQ10*: *qLFI_4* and *qLSI_7*). This locus is particularly interesting because it coincides with the position of *Ghd7*.*1* (approximately 750 kb), a major gene that, in its functional form, delays heading and enhances plant height and yield under long-day conditions (Yan et al., 2013). Moreover, *Ghd7*.*1* has been proposed to influence culm strength (Guo et al, 2021). However, we did not detect a QTL for heading date at the position of *Ghd7*.*1* locus, suggesting that the *PQ8* may not correspond to *Ghd7*.*1* itself. This indicates the possibility that these loci represent new genes that have not been previously reported

In addition to these, we detected moderate-effect QTL on chromosomes 3 and 4 for internode lengths (*PQ4*: *qLFI_1, qLSI_1, qLTI_2* and *qHD_1* on Chr3. *PQ6*: *qLFI_6, qLSI_3* and *qLTI_6* on Chr4). The presence of multiple loci across the genome underscores that internode length is a classic polygenic trait. This concurs with previous studies that have used internode elongation as a proxy for lodging and found QTL dispersed on several chromosomes (Zhu et al, 2008; Yadav et al, 2017; Jiang et al, 2019). It is also noteworthy that C309, the taller parent, produces six elongated internodes as opposed to five in ZS97; this suggests genetic differences in either the initiation of an extra internode or the partitioning of elongation among internodes. Some of the QTL we identified (such as those on chromosome 6 or chromosome 7) might underlie these differences, by controlling the developmental timing or hormonal signals that determine internode number and length distribution along the stem.

Our analysis of internode diameter revealed fewer QTL than for length, but these loci provide important insights into the genetic control of stem strength. We found a single QTL (*PQ11*: *qDFI_7* and *qDSI_5*) on chromosome 8 affecting the diameter of the first and second internode. it did not show a significant correlation with DTI. Although *IPA1/OsSPL14* is a gene known to confer a more ideal plant architecture–including fewer tillers and enhanced culm strength–when an optimal allele is present (Jiao et al, 2010; Chao et al., 2021), the physical distance between this *PQ11* and IPA1 is relatively large (2589.7 kb), suggesting that this locus may represent a novel QTL distinct from *IPA1*. The diameters of the second and third internodes were influenced by overlapping QTL on chromosome 2 (*PQ1*: *qDSI_2* and *qDTI_3* at 45.2 cM. *PQ3*: *qDFI_2* and *qDTI_1* at 137.5 cM) and on chromosome 12 (*PQ14*: *qDSI_7* and *qDTI_2* at 0.7–1.2 cM). Additionally, a unique QTL (*PQ13*: *qLSI_10* and *qLFI_7*) on chromosome 11 specifically affected the first and second internode length. In addition, we identified a QTL (*PQ15*: *qLSI_8* and *qLTI_5*) on chromosome 12 that controls the lengths of the second and third internodes. The clustering of QTL for DSI and DTI on chromosomes 2 and 12 suggests that the genetic control of internode diameter in the lower internodes is partly shared. This is supported by the phenotypic data: we observed an extremely high positive correlation (r = 0.89) between DSI and DTI (Figure S1). *Gn1a* can improve lodging resistance by promoting more crown root growth and culm diameter (Tu et al, 2022). The identified QTL *qLTI_1* is located approximately 339.5 kb from *Gn1a*, raising the possibility that *Gn1a* may also be involved in the regulation of third internode length.

In addition, a minor pleiotropic QTL *PQ9* was noted on chromosome 6 at 111.8 cM, where *qLFI_3, qLSI_6* and *qDFI_6* co-localize, indicating a shared influence on the first and two internodes. It lies near the reported position of *SCM2* (847.1 kb), which encodes the F-box protein gene *APO1* and is known to enhance culm thickness and strength (Ookawa et al, 2010).

### 4.4 Significance of candidate gene selection and haplotype analysis in gene cloning

Candidate genes were selected by analyzing the expression differences in stem tissues between the two parents. However, this approach is considered to have only reference value, as genetic background differences may result in gene interactions that affect gene expression. Therefore, further construction of a backcross population is needed to reduce genetic background interference and enable fine mapping of the target gene. Nevertheless, RNA-Seq was used to significantly narrow down the selection range. Initially, the pleiotropic QTL regions contained 1,263 expressed genes, but after differential gene selection, only 183 genes remained in this study, laying a solid foundation for candidate gene selection. It should be noted that some genes, even if no expression differences are observed, may still exhibit functional differences, as the variation in their function may arise from structural variations.

Additionally, by integrating previously published data and performing haplotype analysis, the impact of different haplotypes on phenotypic traits was accurately assessed, and allele effects were determined. This is crucial for both genetic modification and breeding. For example, Hap3, which was identified, is a haplotype exclusively found in temperate japonica.

### 4.5 Breeding implications and future directions

Improving lodging resistance without compromising yield remains a major challenge in rice breeding, particularly for high-yielding hybrids that are prone to lodging due to increased plant height and heavier panicles. For instance, hybrid rice has been shown to yield 8.9% more than inbred lines but also exhibits significantly higher lodging indices at the basal and third internodes (Liao et al., 2023). Our study identified several QTL that offer molecular targets to address this issue. For example, combining the C309 allele at *qDSI_7/qDTI_2* (Chr12), which enhances lower internode thickness, with the ZS97 allele at *qHD_2* (*Hd1* on Chr6), which moderates heading date and plant height, may produce lines with improved stem strength and reduced lodging risk. These loci provide valuable tools for marker-assisted or genomic selection in breeding programs.

Although two loci controlling culm diameter, *STRONG1* (*LOC_Os06g14080*) and *STRONG2* (*LOC_Os03g26044*), have been cloned recently, neither of them overlaps with the QTLs identified in our study. This suggests that the parental lines used in our population do not carry allelic variations at these loci, which further highlights the genetic diversity present in our mapping population. This observation lays the foundation for future efforts to clone additional QTLs controlling culm-related traits.

Our findings also highlight the value of precise phenotyping and advanced mapping populations in uncovering novel QTL for complex traits. By focusing on internode traits with high heritability and using a dense SNP bin map, we detected loci that were missed in earlier studies. Some of these, such as QTL on chromosomes 2, 7, and 12, warrant further fine-mapping and gene cloning to clarify their roles in culm strength. These discoveries, together with integration of known genes like *Ghd7*.*1, SCM2*, and *Gn1a*, could enable the development of elite varieties with enhanced lodging resistance through pyramiding and gene editing. Ultimately, multi-trait molecular breeding will be essential to achieving yield stability in rice production systems.

## 5 Conclusion

In this study, we identified several quantitative trait loci (QTLs) associated with internode length and diameter, which are critical determinants of lodging resistance in rice. The identification of 54 QTLs, including 15 pleiotropic loci, provides valuable insights into the genetic architecture of these traits. Importantly, some of the QTLs identified were novel, which could be further utilized in marker-assisted selection for breeding rice varieties with improved lodging resistance. Additionally, through RNA-Seq analysis, several candidate genes were identified. Haplotype analysis and LD analysis were performed on *LOC_Os06g11130*, revealing significant haplotype variation and confirming its potential role in lodging resistance. However, this study is limited by the use of a single-year dataset, which may reduce the generalizability of the results. To address this limitation, future studies should incorporate multi-year validation to confirm the stability of these QTLs across different environmental conditions. Additionally, functional follow-up, including fine-mapping and gene cloning, will be crucial to further elucidate the roles of these QTLs and identify candidate genes for enhancing lodging resistance. These efforts will lay the foundation for breeding high-yielding rice varieties with improved lodging resistance, contributing to greater yield stability in the face of environmental stress.

## Data Availability Statement

The next-generation sequencing data used in this study were submitted to the National Genomics Data Center under the BioProject PRJCA044238. Other data presented in this study are available on request from the corresponding author.

## Acknowledge

We would like to express our sincere gratitude to Professor Jianmin Bian from Jiangxi Agricultural University for his valuable guidance and insightful suggestions on this work.

## Author Contributions

W.H. and Z.X. conceived and designed the experiments. Q.L. and W.H. performed most the experiments. P.G., Y,Z., and W.H. analyzed the data. Y.L. conducted field management and plant cultivation. Q.L., P.G. and W.H. wrote the paper. All authors reviewed and approved the final manuscript.

