## Supplementary files for "Identification of QTLs for Internode Length and Diameter Associated with Lodging Resistance in Rice"

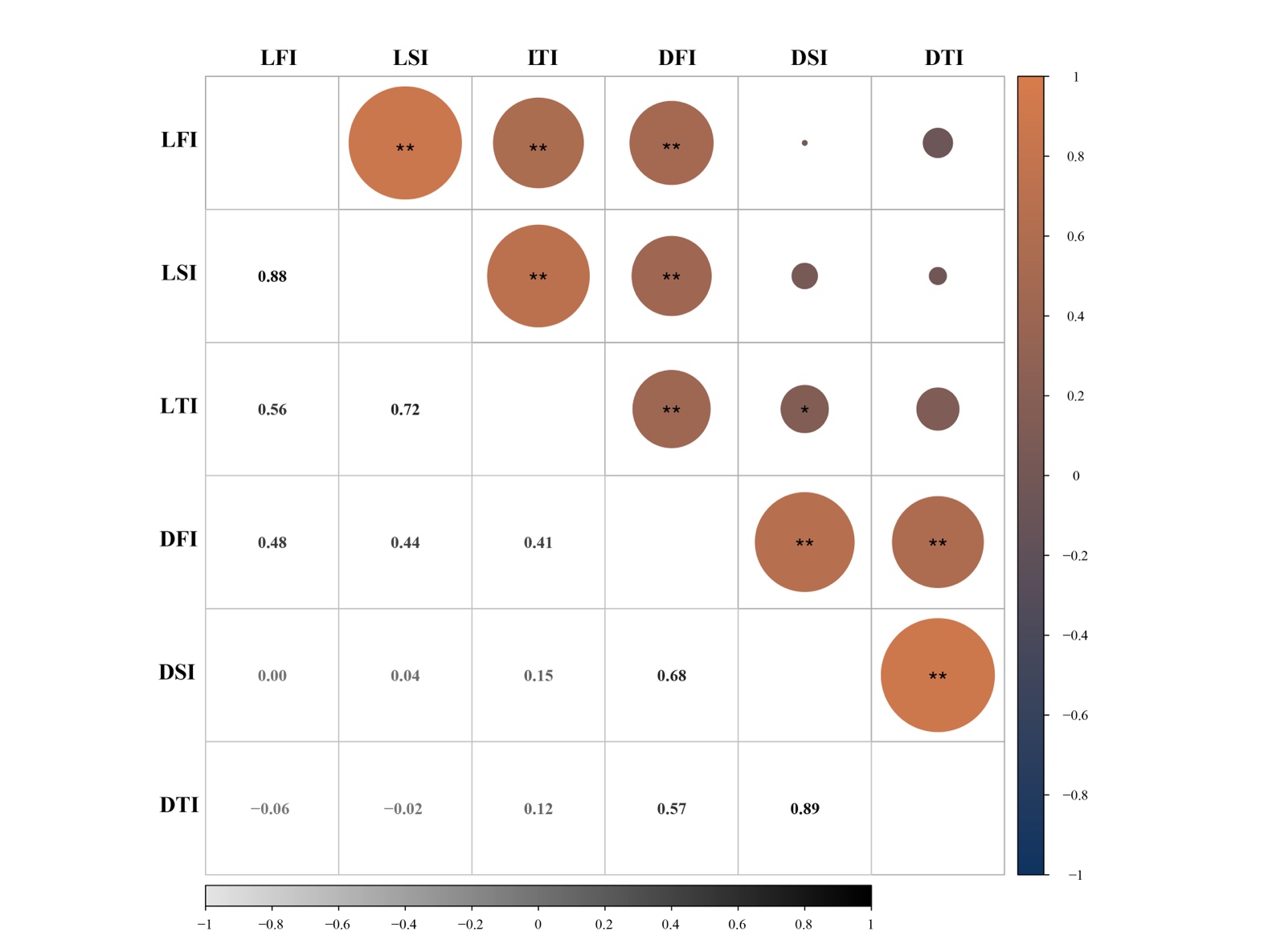

**Figure S1** Correlation between biomass-related traits of single plant in RIL population. * *P*-value < 0.05. ^**^ *P*-value < 0.01.

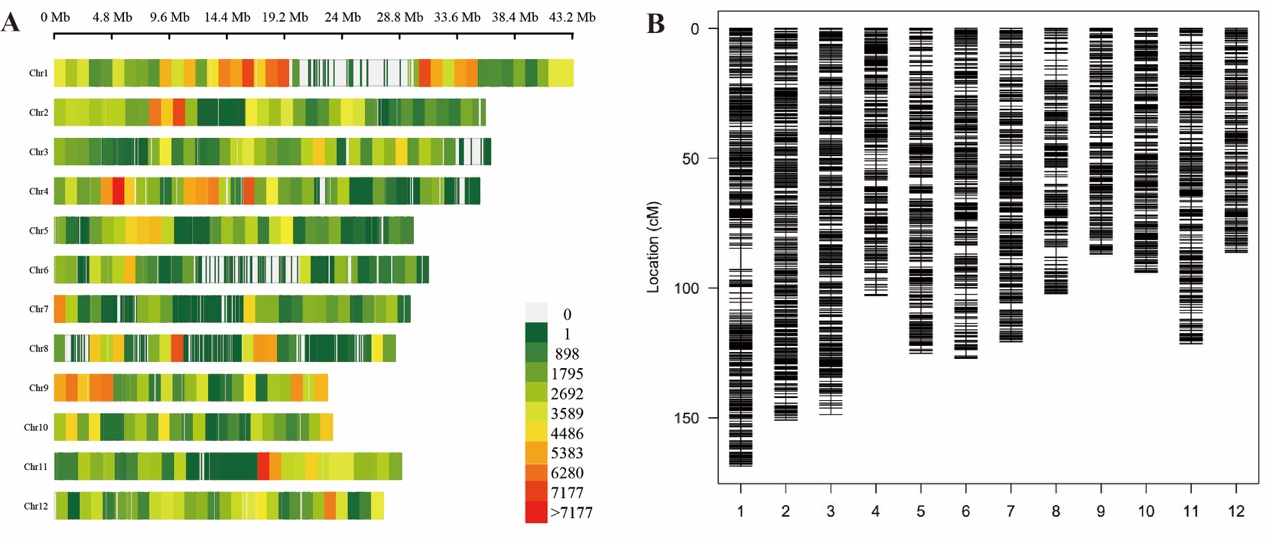

**Figure S2** Genetic analysis of the RIL populations. **(A)**, the SNP density map. **(B)**, the high-density genetic linkage map

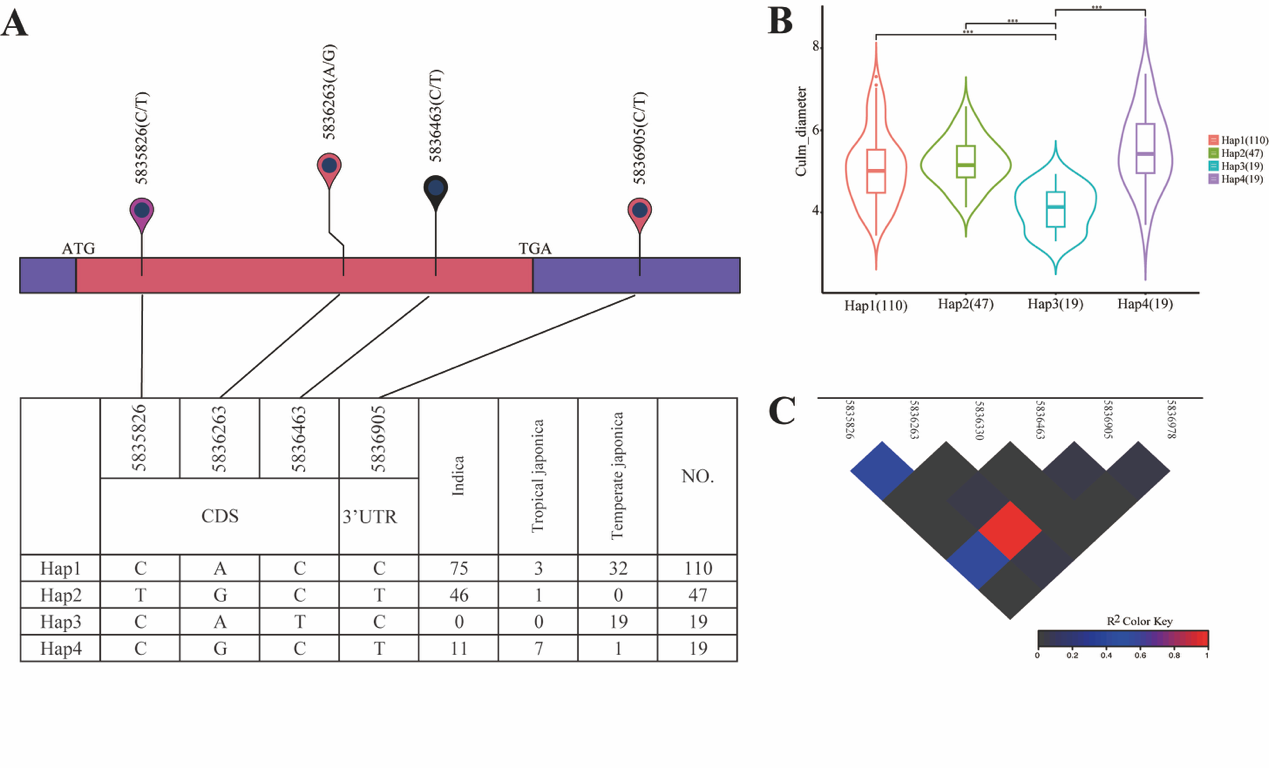

**Figure S3** Natural variation in *LOC_Os06g11130*. **(A)** Haplotypes of *LOC_Os06g11130* in 195 accessions. **(B)** The culm diameter (diameter of first internode in this study) of *LOC_Os06g11130* with different haplotypes. ^***^ *P*-value < 0.001. **(C)**, LD analysis of *LOC_Os06g11130*.

**Table S1** Distribution of BINs on chromosomes.

| Chromosomes | Number of BINs | Length (cM) | Average distance (cM) | Maximum gap (cM) |
| --- | --- | --- | --- | --- |
| 1 | 374 | 168.6 | 0.45 | 8.2 |
| 2 | 375 | 150.9 | 0.40 | 2.0 |
| 3 | 361 | 148.7 | 0.41 | 2.5 |
| 4 | 242 | 102.8 | 0.43 | 2.8 |
| 5 | 284 | 125.0 | 0.44 | 3.1 |
| 6 | 261 | 127.1 | 0.49 | 3.6 |
| 7 | 270 | 120.8 | 0.45 | 2.8 |
| 8 | 203 | 102.3 | 0.51 | 4.2 |
| 9 | 215 | 86.9 | 0.41 | 2.3 |
| 10 | 235 | 93.8 | 0.40 | 1.8 |
| 11 | 286 | 121.6 | 0.43 | 2.6 |
| 12 | 201 | 86.3 | 0.43 | 1.7 |
| Average | 275.6 | 119.6 | 0.44 | 3.1 |
| Total | 3307 | 1434.7 | - | - |

**Table S2** QTL identified for all the traits.

| QTLs | Environments | Chr | Position (cM) | Additive Effect | LOD | Left_Marker | Right_Marker | Var_Genet_(i) | PVE (%) | Var_Error | Var_Phen (total) |
| --- | --- | --- | --- | --- | --- | --- | --- | --- | --- | --- | --- |
| *qHD_1* | R1 | 3 | 3.4 | -2.6 | 8.0 | Chr3_1165038 | Chr3_1165038 | 6.8 | 12.3 | 31.0 | 55.4 |
| *qHD_2* | R1 | 6 | 49.9 | 2.8 | 9.6 | Chr6_9256490 | Chr6_9256490 | 8.0 | 14.5 |  |  |
| *qHD_3* | R1 | 7 | 56.8 | 1.9 | 4.5 | Chr7_10064656 | Chr7_10105892 | 3.8 | 6.8 |  |  |
| *qHD_4* | R1 | 10 | 79.5 | 2.4 | 6.4 | Chr10_20763344 | Chr10_20763344 | 5.7 | 10.3 |  |  |
| *qHD_1* | R2 | 3 | 3.4 | -2.7 | 10.7 | Chr3_1165038 | Chr3_1165038 | 7.3 | 16.3 | 24.9 | 45.1 |
| *qHD_2* | R2 | 6 | 46.1 | 2.4 | 8.6 | Chr6_7910387 | Chr6_7910387 | 5.6 | 12.4 |  |  |
| *qHD_3* | R2 | 7 | 56.7 | 1.8 | 4.8 | Chr7_10064656 | Chr7_10064656 | 3.2 | 7.0 |  |  |
| *qHD_4* | R2 | 10 | 79.5 | 2.0 | 5.9 | Chr10_20763344 | Chr10_20763344 | 4.2 | 9.2 |  |  |
| *qHD_1* | R3 | 3 | 3.4 | -2.6 | 10.0 | Chr3_1165038 | Chr3_1165038 | 6.8 | 15.2 | 25.3 | 45.1 |
| *qHD_2* | R3 | 6 | 46.6 | 2.3 | 8.2 | Chr6_7962320 | Chr6_7962320 | 5.4 | 12.0 |  |  |
| *qHD_3* | R3 | 7 | 56.7 | 1.8 | 5.0 | Chr7_10064656 | Chr7_10064656 | 3.3 | 7.4 |  |  |
| *qHD_4* | R3 | 10 | 79.5 | 2.1 | 5.9 | Chr10_20763344 | Chr10_20763344 | 4.2 | 9.4 |  |  |
| *qLFI_1* | R1 | 3 | 2.7 | -0.3 | 3.1 | Chr3_991280 | Chr3_991280 | 0.1 | 5.7 | 1.0 | 1.6 |
| *qLFI_2* | R1 | 6 | 35.8 | 0.5 | 6.6 | Chr6_6577509 | Chr6_6577509 | 0.2 | 13.7 |  |  |
| *qLFI_3* | R1 | 6 | 111.8 | 0.3 | 4.1 | Chr6_28052937 | Chr6_28052937 | 0.1 | 7.2 |  |  |
| *qLFI_4* | R1 | 7 | 118.9 | 0.4 | 4.1 | Chr7_28864269 | Chr7_28864269 | 0.1 | 9.1 |  |  |
| *qLFI_5* | R2 | 6 | 17.2 | 0.4 | 3.4 | Chr6_3117614 | Chr6_3117614 | 0.1 | 8.0 | 1.5 | 1.8 |
| *qLFI_3* | R2 | 6 | 111.8 | 0.4 | 3.4 | Chr6_28052937 | Chr6_28052937 | 0.1 | 8.0 |  |  |
| *qLFI_1* | R3 | 3 | 3.9 | -0.5 | 6.0 | Chr3_1229922 | Chr3_1229922 | 0.2 | 9.6 | 1.2 | 2.4 |
| *qLFI_6* | R3 | 4 | 13.6 | -0.4 | 4.2 | Chr4_11866366 | Chr4_11866366 | 0.2 | 7.6 |  |  |
| *qLFI_2* | R3 | 6 | 34.1 | 0.5 | 6.8 | Chr6_5879246 | Chr6_5879246 | 0.3 | 11.3 |  |  |
| *qLFI_4* | R3 | 7 | 119.4 | 0.5 | 6.0 | Chr7_28878316 | Chr7_28878316 | 0.3 | 10.9 |  |  |
| *qLFI_7* | R3 | 11 | 64.3 | 0.5 | 6.6 | Chr11_16453511 | Chr11_16453511 | 0.2 | 10.3 |  |  |
| *qLSI_1* | R1 | 3 | 3.4 | -0.6 | 4.7 | Chr3_1165038 | Chr3_1165038 | 0.3 | 6.1 | 3.7 | 5.7 |
| *qLSI_2* | R1 | 3 | 128.9 | -0.4 | 3.0 | Chr3_30774855 | Chr3_30774855 | 0.2 | 2.9 |  |  |
| *qLSI_3* | R1 | 4 | 10.7 | -0.5 | 3.2 | Chr4_5318807 | Chr4_5318807 | 0.3 | 4.5 |  |  |
| *qLSI_4* | R1 | 5 | 10.5 | 0.4 | 3.0 | Chr5_1690358 | Chr5_1690358 | 0.2 | 2.9 |  |  |
| *qLSI_5* | R1 | 6 | 35.8 | 0.7 | 5.7 | Chr6_6577509 | Chr6_6577509 | 0.4 | 7.9 |  |  |
| *qLSI_6* | R1 | 6 | 111.8 | 0.5 | 3.9 | Chr6_28052937 | Chr6_28052937 | 0.3 | 4.6 |  |  |
| *qLSI_7* | R1 | 7 | 118.9 | 0.5 | 2.7 | Chr7_28864269 | Chr7_28864269 | 0.2 | 4.0 |  |  |
| *qLSI_8* | R1 | 12 | 19.8 | 0.4 | 2.9 | Chr12_3359883 | Chr12_3359883 | 0.2 | 2.8 |  |  |
| *qLSI_3* | R2 | 4 | 9.4 | -0.6 | 2.7 | Chr4_5291486 | Chr4_5291486 | 0.3 | 6.8 | 3.7 | 4.7 |
| *qLSI_9* | R2 | 6 | 16.7 | 0.5 | 2.7 | Chr6_2980606 | Chr6_2980606 | 0.3 | 6.0 |  |  |
| *qLSI_6* | R2 | 6 | 111.8 | 0.6 | 3.6 | Chr6_28052937 | Chr6_28052937 | 0.4 | 8.2 |  |  |
| *qLSI_1* | R3 | 3 | 2.7 | -0.8 | 5.9 | Chr3_991280 | Chr3_991280 | 0.6 | 11.0 | 3.6 | 5.8 |
| *qLSI_5* | R3 | 6 | 35.3 | 0.9 | 6.9 | Chr6_6252782 | Chr6_6252782 | 0.8 | 13.2 |  |  |
| *qLSI_10* | R3 | 11 | 61.6 | 0.7 | 4.9 | Chr11_10951616 | Chr11_10951616 | 0.5 | 8.6 |  |  |
| *qLSI_8* | R3 | 12 | 19.8 | 0.6 | 3.1 | Chr12_3359883 | Chr12_3359883 | 0.3 | 5.4 |  |  |
| *qLTI_1* | R1 | 1 | 27.5 | 0.7 | 2.7 | Chr1_4930628 | Chr1_4930628 | 0.4 | 5.0 | 6.4 | 8.6 |
| *qLTI_2* | R1 | 3 | 3.4 | -0.7 | 3.5 | Chr3_1165038 | Chr3_1165038 | 0.5 | 5.5 |  |  |
| *qLTI_3* | R1 | 3 | 85.5 | -0.6 | 2.7 | Chr3_18437518 | Chr3_18437518 | 0.4 | 4.2 |  |  |
| *qLTI_4* | R1 | 6 | 34.1 | 0.7 | 3.3 | Chr6_5879246 | Chr6_5879246 | 0.5 | 5.9 |  |  |
| *qLTI_5* | R1 | 12 | 19.8 | 0.6 | 2.5 | Chr12_3359883 | Chr12_3359883 | 0.4 | 4.4 |  |  |
| *qLTI_2* | R2 | 3 | 6.1 | -0.9 | 3.1 | Chr3_1980534 | Chr3_1980534 | 0.7 | 6.1 | 9.1 | 12.3 |
| *qLTI_6* | R2 | 4 | 13.8 | -0.9 | 3.2 | Chr4_12407413 | Chr4_12407413 | 0.8 | 6.5 |  |  |
| *qLTI_7* | R2 | 11 | 64.3 | 1.0 | 4.1 | Chr11_16453511 | Chr11_16453511 | 0.9 | 7.4 |  |  |
| *qLTI_5* | R2 | 12 | 19.8 | 0.9 | 3.3 | Chr12_3359883 | Chr12_3359883 | 0.8 | 6.4 |  |  |
| *qLTI_3* | R3 | 3 | 90.1 | -0.7 | 3.0 | Chr3_23481393 | Chr3_23481393 | 0.5 | 6.2 | 7.0 | 8.6 |
| *qLTI_4* | R3 | 6 | 34.1 | 0.8 | 3.9 | Chr6_5879246 | Chr6_5879246 | 0.7 | 7.8 |  |  |
| *qLTI_5* | R3 | 12 | 19.8 | 0.7 | 2.7 | Chr12_3359883 | Chr12_3359883 | 0.5 | 5.5 |  |  |
| *qDFI_1* | R1 | 1 | 72.5 | 0.2 | 2.8 | Chr1_18454238 | Chr1_18454238 | 0.0 | 4.7 | 0.4 | 0.6 |
| *qDFI_2* | R1 | 2 | 137.5 | -0.2 | 3.0 | Chr2_33989267 | Chr2_33989267 | 0.0 | 4.4 |  |  |
| *qDFI_3* | R1 | 4 | 75.1 | -0.1 | 3.0 | Chr4_27435819 | Chr4_27435819 | 0.0 | 4.0 |  |  |
| *qDFI_4* | R1 | 5 | 77.6 | -0.1 | 2.9 | Chr5_20697160 | Chr5_20697160 | 0.0 | 3.5 |  |  |
| *qDFI_5* | R1 | 6 | 30.7 | 0.2 | 4.1 | Chr6_5205005 | Chr6_5205005 | 0.0 | 7.7 |  |  |
| *qDFI_6* | R1 | 6 | 118.5 | 0.2 | 3.5 | Chr6_29068108 | Chr6_29068108 | 0.0 | 6.5 |  |  |
| *qDFI_5* | R2 | 6 | 30.7 | 0.2 | 3.1 | Chr6_5205005 | Chr6_5205005 | 0.0 | 7.4 | 0.3 | 0.4 |
| *qDFI_7* | R2 | 8 | 102.3 | -0.2 | 3.1 | Chr8_27868446 | Chr8_27868446 | 0.0 | 7.4 |  |  |
| *qDFI_8* | R3 | 2 | 1.0 | -0.2 | 2.8 | Chr2_1023542 | Chr2_1023542 | 0.0 | 5.3 | 0.4 | 0.6 |
| *qDFI_5* | R3 | 6 | 30.7 | 0.2 | 3.4 | Chr6_5205005 | Chr6_5205005 | 0.0 | 6.5 |  |  |
| *qDFI_6* | R3 | 6 | 114.5 | 0.2 | 3.9 | Chr6_28327261 | Chr6_28327261 | 0.0 | 6.6 |  |  |
| *qDFI_9* | R3 | 9 | 34.3 | -0.2 | 2.7 | Chr9_12987140 | Chr9_12987140 | 0.0 | 5.1 |  |  |
| *qDSI_1* | R1 | 5 | 38.0 | -0.2 | 2.8 | Chr5_5821403 | Chr5_5821403 | 0.0 | 6.9 | 0.3 | 0.3 |
| *qDSI_2* | R2 | 1 | 45.2 | 0.2 | 3.2 | Chr1_8270335 | Chr1_8270335 | 0.0 | 4.6 | 0.4 | 0.6 |
| *qDSI_3* | R2 | 2 | 123.4 | -0.2 | 3.2 | Chr2_30724778 | Chr2_30724778 | 0.0 | 6.2 |  |  |
| *qDSI_4* | R2 | 3 | 75.5 | -0.2 | 2.9 | Chr3_15829265 | Chr3_15829265 | 0.0 | 4.0 |  |  |
| *qDSI_1* | R2 | 5 | 39.0 | -0.2 | 3.1 | Chr5_6247006 | Chr5_6247006 | 0.0 | 4.7 |  |  |
| *qDSI_5* | R2 | 8 | 102.3 | -0.2 | 2.9 | Chr8_27868446 | Chr8_27868446 | 0.0 | 4.1 |  |  |
| *qDSI_6* | R2 | 9 | 83.8 | -0.2 | 2.7 | Chr9_21352300 | Chr9_21352300 | 0.0 | 4.4 |  |  |
| *qDSI_1* | R3 | 5 | 36.8 | -0.1 | 2.6 | Chr5_5458989 | Chr5_5458989 | 0.0 | 6.1 | 0.3 | 0.3 |
| *qDSI_7* | R3 | 12 | 0.7 | 0.1 | 3.0 | Chr12_703295 | Chr12_703295 | 0.0 | 7.1 |  |  |
| *qDTI_1* | R1 | 2 | 137.5 | -0.2 | 3.5 | Chr2_33989267 | Chr2_33989267 | 0.0 | 7.7 | 0.3 | 0.3 |
| *qDTI_2* | R1 | 12 | 0.7 | 0.2 | 4.8 | Chr12_703295 | Chr12_703295 | 0.0 | 10.8 |  |  |
| *qDTI_3* | R2 | 1 | 45.2 | 0.2 | 5.1 | Chr1_8270335 | Chr1_8270335 | 0.0 | 7.1 | 0.4 | 0.6 |
| *qDTI_4* | R2 | 2 | 123.4 | -0.2 | 3.7 | Chr2_30724778 | Chr2_30724778 | 0.0 | 7.2 |  |  |
| *qDTI_5* | R2 | 3 | 75.5 | -0.2 | 3.0 | Chr3_15829265 | Chr3_15829265 | 0.0 | 4.2 |  |  |
| *qDTI_6* | R2 | 9 | 83.3 | -0.2 | 3.6 | Chr9_21319920 | Chr9_21319920 | 0.0 | 4.1 |  |  |
| *qDTI_7* | R2 | 10 | 5.4 | -0.2 | 2.9 | Chr10_1661619 | Chr10_1661619 | 0.0 | 4.4 |  |  |
| *qDTI_8* | R3 | 2 | 101.3 | -0.2 | 2.8 | Chr2_24226745 | Chr2_24226745 | 0.0 | 4.4 |  |  |
| *qDTI_9* | R3 | 7 | 88.7 | -0.2 | 3.7 | Chr7_24487072 | Chr7_24487072 | 0.0 | 7.1 | 0.4 | 0.6 |
| *qDTI_10* | R3 | 11 | 29.3 | 0.2 | 4.0 | Chr11_3846236 | Chr11_3846236 | 0.0 | 6.5 |  |  |
| *qDTI_2* | R3 | 12 | 1.2 | 0.2 | 3.8 | Chr12_734214 | Chr12_734214 | 0.0 | 6.4 |  |  |

Note: HD, heading date. LFI, length of the first internode. LSI, length of the second internode. LTI, length of the third internode. DFI, diameter of the first internode. DSI, diameter of the second internode. DTI, diameter of the third internode. Chr, chromosome. LOD, likelihood of odd. PVE, phenotypic variance explained. Var_Genet_(i) indicates the genetic variance explained by the QTL. Var_Error represents the error variance. Var_Phen (total) denotes the total phenotypic variance. R1, R2, and R3 represent the three field environments (Nada, Wangwu, and Dongcheng, respectively).

**Table S3** The distribution of pleiotropic QTLs on chromosomes.

| Name of pleiotropic QTLs | Chr | Interval (cM) | Left_Marker | Right_Marker | Number of genes within the Interval | Number of differentially expressed genes | QTL name |
| --- | --- | --- | --- | --- | --- | --- | --- |
| *PQ1* | 1 | 45.2 | Chr1_8270335 | Chr1_8270335 | 1 | 0 | *qDSI_2* and *qDTI_3* |
| *PQ2* | 2 | 123.4 | Chr2_30724778 | Chr2_30724778 | 1 | 1 | *qDSI_3* and *qDTI_4* |
| *PQ3* | 2 | 137.5 | Chr2_33989267 | Chr2_33989267 | 2 | 0 | *qDFI_2* and *qDTI_1* |
| *PQ4* | 3 | 2.7–6.1 | Chr3_991280 | Chr3_1980534 | 134 | 27 | *qLFI_1*, *qLSI_1*, *qLTI_2* and *qHD_1* |
| *PQ5* | 3 | 75.5 | Chr3_15829265 | Chr3_15829265 | 2 | 0 | *qDSI_4* and *qDTI_5* |
| *PQ6* | 4 | 9.4–13.8 | Chr4_5291486 | Chr4_12407413 | 436 | 64 | *qLFI_6*, *qLSI_3* and *qLTI_6* |
| *PQ7* | 6 | 16.7–17.2 | Chr6_2980606 | Chr6_3117614 | 24 | 4 | *qLSI_9* and *qLFI_5* |
| *PQ8* | 6 | 30.7–35.8 | Chr6_5205005 | Chr6_6577509 | 178 | 26 | *qDFI_5*, *qLFI_2*, *qLTI_4* and *qLSI_5* |
| *PQ9* | 6 | 111.8–118.5 | Chr6_28052937 | Chr6_29068108 | 150 | 16 | *qLFI_3*, *qLSI_6* and *qDFI_6* |
| *PQ10* | 7 | 118.9–119.4 | Chr7_28864269 | Chr7_28878316 | 2 | 0 | *qLFI_4* and *qLSI_7* |
| *PQ11* | 8 | 102.3 | Chr8_27868446 | Chr8_27868446 | 2 | 0 | *qDFI_7* and *qDSI_5* |
| *PQ12* | 9 | 83.3–83.8 | Chr9_21319920 | Chr9_21352300 | 9 | 0 | *qDTI_6* and *qDSI_6* |
| *PQ13* | 11 | 61.6–64.3 | Chr11_10951616 | Chr11_16453511 | 314 | 43 | *qLSI_10* and *qLFI_7* |
| *PQ14* | 12 | 0.7–1.2 | Chr12_703295 | Chr12_734214 | 6 | 2 | *qDSI_7* and *qDTI_2* |
| *PQ15* | 12 | 19.8 | Chr12_3359883 | Chr12_3359883 | 2 | 0 | *qLSI_8* and *qLTI_5* |

**Table S4** Differentially Expressed Genes and Annotation Information within pleiotropic -QTL Regions.

| Name of pleiotropic QTLs | Gene ID | Nipponbare_Stem  (log2(TPM + 1)) | Zhenshan 97_Stem  (log2(TPM + 1)) | \|log2FC\| | annotation |
| --- | --- | --- | --- | --- | --- |
| *PQ2* | *LOC_Os02g50320* | 2.25 | 3.3 | 1.05 | expressed protein |
| *PQ4* | *LOC_Os03g02710* | 2.59 | 1.45 | 1.14 | hydroxymethylglutaryl-CoA synthase, putative, expressed |
|  | *LOC_Os03g02780* | 3.22 | 1.94 | 1.28 | Core histone H2A/H2B/H3/H4 domain containing protein, putative, expressed |
|  | *LOC_Os03g02800* | 0.45 | 2.25 | 1.8 | no apical meristem protein, putative, expressed |
|  | *LOC_Os03g02840* | 3.41 | 4.47 | 1.06 | remorin family protein, putative, expressed |
|  | *LOC_Os03g02920* | 2.45 | 3.96 | 1.51 | peroxidase precursor, putative, expressed |
|  | *LOC_Os03g02960* | 5.75 | 4.25 | 1.5 | nascent polypeptide-associated complex subunit alpha, putative, expressed |
|  | *LOC_Os03g03020* | 6.58 | 5 | 1.58 | L11 domain containing ribosomal protein, putative, expressed |
|  | *LOC_Os03g03164* | 3.11 | 5.42 | 2.31 | homeobox protein knotted-1, putative, expressed |
|  | *LOC_Os03g03200* | 3.16 | 4.43 | 1.27 | hydrolase, alpha/beta fold family protein, putative, expressed |
|  | *LOC_Os03g03360* | 6.01 | 4.93 | 1.08 | ribosomal protein L5, putative, expressed |
|  | *LOC_Os03g03370* | 1.23 | 2.54 | 1.31 | fatty acid hydroxylase, putative, expressed |
|  | *LOC_Os03g03460* | 6.8 | 5.61 | 1.19 | erythronate-4-phosphate dehydrogenase, putative, expressed |
|  | *LOC_Os03g03470* | 3.98 | 2.07 | 1.91 | expressed protein |
|  | *LOC_Os03g03490* | 4.71 | 0 | 4.71 | expressed protein |
|  | *LOC_Os03g03500* | 5.1 | 2.7 | 2.4 | heavy metal-associated domain containing protein, expressed |
|  | *LOC_Os03g03570* | 0.07 | 1.15 | 1.08 | leucine-rich repeat transmembrane protein kinase, putative, expressed |
|  | *LOC_Os03g03590* | 3.45 | 4.74 | 1.29 | transporter, monovalent cation:proton antiporter-2 family, putative, expressed |
|  | *LOC_Os03g03720* | 7.47 | 5.62 | 1.85 | glyceraldehyde-3-phosphate dehydrogenase, putative, expressed |
|  | *LOC_Os03g03724* | 11.35 | 8.79 | 2.56 | expressed protein |
|  | *LOC_Os03g03730* | 0.48 | 1.94 | 1.46 | regulatory protein, putative, expressed |
|  | *LOC_Os03g03760* | 0.18 | 1.28 | 1.1 | MYB family transcription factor, putative, expressed |
|  | *LOC_Os03g03910* | 2.38 | 0.94 | 1.44 | catalase domain containing protein, expressed |
|  | *LOC_Os03g03990* | 3.36 | 1.56 | 1.8 | signal recognition particle 43 kDa protein, chloroplast precursor, putative, expressed |
|  | *LOC_Os03g04130* | 4 | 1.93 | 2.07 | AMP-binding domain containing protein, expressed |
|  | *LOC_Os03g04220* | 0.82 | 4.64 | 3.82 | glutathione S-transferase, putative, expressed |
|  | *LOC_Os03g04240* | 0.44 | 5.86 | 5.42 | glutathione S-transferase, putative, expressed |
|  | *LOC_Os03g04250* | 0.23 | 3.99 | 3.76 | glutathione S-transferase, putative, expressed |
| *PQ6* | *LOC_Os04g10214* | 1.26 | 0 | 1.26 | expressed protein |
|  | *LOC_Os04g10460* | 4.19 | 2.86 | 1.33 | amidase, putative, expressed |
|  | *LOC_Os04g10760* | 0 | 1.7 | 1.7 | expressed protein |
|  | *LOC_Os04g11120* | 1.27 | 0 | 1.27 | expressed protein |
|  | *LOC_Os04g11390* | 0.62 | 3 | 2.38 | expressed protein |
|  | *LOC_Os04g11840* | 3.95 | 1.79 | 2.16 | expressed protein |
|  | *LOC_Os04g11890* | 1 | 2.11 | 1.11 | OsFBX121 - F-box domain containing protein, expressed |
|  | *LOC_Os04g11980* | 3.85 | 0 | 3.85 | expressed protein |
|  | *LOC_Os04g12140* | 1.97 | 0 | 1.97 | expressed protein |
|  | *LOC_Os04g12460* | 3.73 | 0 | 3.73 | Leucine Rich Repeat family protein, expressed |
|  | *LOC_Os04g12499* | 0.24 | 5.9 | 5.66 | amino acid transporter protein, putative, expressed |
|  | *LOC_Os04g12530* | 0.24 | 1.29 | 1.05 | amino acid transporter family protein, putative, expressed |
|  | *LOC_Os04g12600* | 2.31 | 0 | 2.31 | receptor-like protein kinase, putative, expressed |
|  | *LOC_Os04g12744* | 3.49 | 0 | 3.49 | expressed protein |
|  | *LOC_Os04g12820* | 3.23 | 0.99 | 2.24 | expressed protein |
|  | *LOC_Os04g12900* | 0 | 2.97 | 2.97 | glucosyltransferase, putative, expressed |
|  | *LOC_Os04g12980* | 5.14 | 2.34 | 2.8 | UDP-glucoronosyl/UDP-glucosyl transferase, putative, expressed |
|  | *LOC_Os04g13140* | 1.59 | 0.43 | 1.16 | vignain precursor, putative, expressed |
|  | *LOC_Os04g13150* | 1.93 | 0.19 | 1.74 | OsFBX125 - F-box domain containing protein, expressed |
|  | *LOC_Os04g13170* | 1.38 | 0.33 | 1.05 | OsFBD10 - F-box and FBD domain containing protein, expressed |
|  | *LOC_Os04g13260* | 0.18 | 3.97 | 3.79 | expressed protein |
|  | *LOC_Os04g14110* | 3.53 | 0.08 | 3.45 | TRAF-type zinc finger family protein, expressed |
|  | *LOC_Os04g14150* | 3.52 | 5.37 | 1.85 | dehydration response related protein, putative, expressed |
|  | *LOC_Os04g14200* | 2.19 | 0 | 2.19 | expressed protein |
|  | *LOC_Os04g14204* | 3.29 | 0.14 | 3.15 | expressed protein |
|  | *LOC_Os04g14220* | 2.5 | 0 | 2.5 | disease resistance protein RPM1, putative, expressed |
|  | *LOC_Os04g14304* | 2.63 | 0 | 2.63 | OsWAK30/31 - OsWAK receptor-like protein OsWAK-RLP, expressed |
|  | *LOC_Os04g14410* | 2.03 | 3.08 | 1.05 | expressed protein |
|  | *LOC_Os04g14450* | 3.05 | 4.35 | 1.3 | pentatricopeptide, putative, expressed |
|  | *LOC_Os04g14680* | 3.99 | 5.4 | 1.41 | OsAPx3 - Peroxisomal Ascorbate Peroxidase encoding gene 5,8, expressed |
|  | *LOC_Os04g14690* | 5.7 | 2.23 | 3.47 | flavin-containing monooxygenase family protein, putative, expressed |
|  | *LOC_Os04g14710* | 2.58 | 0.04 | 2.54 | flavin-containing monooxygenase family protein, putative, expressed |
|  | *LOC_Os04g14730* | 2.86 | 0 | 2.86 | expressed protein |
|  | *LOC_Os04g14810* | 0.96 | 3.2 | 2.24 | 3-5 exonuclease domain-containing protein, putative, expressed |
|  | *LOC_Os04g15550* | 4.6 | 0.06 | 4.54 | expressed protein |
|  | *LOC_Os04g15660* | 3.72 | 0.1 | 3.62 | receptor kinase, putative, expressed |
|  | *LOC_Os04g15670* | 2.92 | 1.68 | 1.24 | expressed protein |
|  | *LOC_Os04g15820* | 2.58 | 3.61 | 1.03 | expressed protein |
|  | *LOC_Os04g15920* | 2.89 | 1.79 | 1.1 | dehydrogenase, putative, expressed |
|  | *LOC_Os04g16680* | 5.81 | 3.58 | 2.23 | fructose-1,6-bisphosphatase, putative, expressed |
|  | *LOC_Os04g16722* | 6.9 | 5.86 | 1.04 | uncharacterized protein ycf68, putative, expressed |
|  | *LOC_Os04g16740* | 4.63 | 3.54 | 1.09 | ATP synthase subunit alpha, putative, expressed |
|  | *LOC_Os04g16748* | 3.99 | 2.41 | 1.58 | ATP synthase B chain, putative, expressed |
|  | *LOC_Os04g16770* | 1.6 | 2.86 | 1.26 | photosynthetic reaction center protein, putative, expressed |
|  | *LOC_Os04g16775* | 4.68 | 2.93 | 1.75 | conserved hypothetical protein |
|  | *LOC_Os04g16782* | 2.08 | 0.45 | 1.63 | chloroplast 50S ribosomal protein L23, putative |
|  | *LOC_Os04g16812* | 2.08 | 0.45 | 1.63 | chloroplast 50S ribosomal protein L23, putative |
|  | *LOC_Os04g16844* | 0 | 1.14 | 1.14 | cytochrome b6-f complex subunit 4, putative, expressed |
|  | *LOC_Os04g16872* | 4.1 | 2.9 | 1.2 | photosystem II D2 protein, putative, expressed |
|  | *LOC_Os04g16874* | 3.88 | 2.05 | 1.83 | photosystem II 44 kDa reaction center protein, putative, expressed |
|  | *LOC_Os04g17050* | 5.24 | 3.82 | 1.42 | glutaredoxin family protein, expressed |
|  | *LOC_Os04g17479* | 4.22 | 0.16 | 4.06 | expressed protein |
|  | *LOC_Os04g17650* | 0.53 | 3.95 | 3.42 | sucrose synthase, putative, expressed |
|  | *LOC_Os04g18030* | 1.62 | 2.64 | 1.02 | Sec1 family transport protein, putative, expressed |
|  | *LOC_Os04g18090* | 8.9 | 7.82 | 1.08 | histone H1, putative, expressed |
|  | *LOC_Os04g20210* | 3.46 | 0.29 | 3.17 | expressed protein |
|  | *LOC_Os04g20270* | 4.57 | 5.82 | 1.25 | transcriptional regulator Sir2 family protein, putative, expressed |
|  | *LOC_Os04g20280* | 1.81 | 4.03 | 2.22 | expressed protein |
|  | *LOC_Os04g20590* | 4.11 | 5.2 | 1.09 | YL1 nuclear protein C-terminal domain containing protein, expressed |
|  | *LOC_Os04g21130* | 0 | 1.12 | 1.12 | F-box protein PP2-B1, putative, expressed |
|  | *LOC_Os04g21320* | 3.31 | 4.69 | 1.38 | membrane associated DUF588 domain containing protein, putative, expressed |
|  | *LOC_Os04g21600* | 2.6 | 4.25 | 1.65 | expressed protein |
|  | *LOC_Os04g21840* | 2.13 | 0 | 2.13 | expressed protein |
|  | *LOC_Os04g21850* | 5.39 | 0 | 5.39 | expressed protein |
| *PQ7* | *LOC_Os06g06490* | 1.71 | 0.29 | 1.42 | U-box domain containing heat shock protein, putative, expressed |
|  | *LOC_Os06g06550* | 0.67 | 4.17 | 3.5 | plant protein of unknown function domain containing protein, expressed |
|  | *LOC_Os06g06580* | 0 | 3.65 | 3.65 | expressed protein |
|  | *LOC_Os06g06590* | 0 | 1.67 | 1.67 | expressed protein |
| *PQ8* | *LOC_Os06g10170* | 1.91 | 2.94 | 1.03 | flavin-containing monooxygenase family protein, putative, expressed |
|  | *LOC_Os06g10230* | 2.46 | 3.83 | 1.37 | receptor-like protein kinase 5 precursor, putative, expressed |
|  | *LOC_Os06g10280* | 3.69 | 4.86 | 1.17 | CDP-alcohol phosphatidyltransferase, putative, expressed |
|  | *LOC_Os06g10355* | 1.12 | 0 | 1.12 | expressed protein |
|  | *LOC_Os06g10410* | 2.97 | 1.77 | 1.2 | uncharacterized mscS family protein, putative, expressed |
|  | *LOC_Os06g10420* | 1.79 | 0.36 | 1.43 | nitrilase, putative, expressed |
|  | *LOC_Os06g10480* | 2.92 | 0.78 | 2.14 | expressed protein |
|  | *LOC_Os06g10510* | 1.4 | 0.06 | 1.34 | oxidoreductase/ transition metal ion |
|  | *LOC_Os06g10560* | 2.11 | 3.14 | 1.03 | leaf senescence related protein, putative, expressed |
|  | *LOC_Os06g10670* | 1.46 | 4.25 | 2.79 | aspartic proteinase nepenthesin-1 precursor, putative, expressed |
|  | *LOC_Os06g10840* | 2.4 | 0 | 2.4 | expressed protein |
|  | *LOC_Os06g10880* | 2.7 | 1.55 | 1.15 | bZIP transcription factor, putative, expressed |
|  | *LOC_Os06g10890* | 6.38 | 8.69 | 2.31 | sterol carrier protein-2, putative, expressed |
|  | *LOC_Os06g11130* | 3.24 | 5.2 | 1.96 | gibberellin receptor GID1L2, putative, expressed |
|  | *LOC_Os06g11135* | 5.44 | 6.44 | 1 | gibberellin receptor GID1L2, putative, expressed |
|  | *LOC_Os06g11310* | 2.97 | 5.11 | 2.14 | plastocyanin-like domain containing protein, putative, expressed |
|  | *LOC_Os06g11380* | 3.6 | 1.98 | 1.62 | geminivirus Rep-interacting motor protein, putative, expressed |
|  | *LOC_Os06g11490* | 2.97 | 7.33 | 4.36 | plastocyanin-like domain containing protein, putative, expressed |
|  | *LOC_Os06g11500* | 3.98 | 1.75 | 2.23 | MCM9 - Putative minichromosome maintenance MCM family subunit 9, expressed |
|  | *LOC_Os06g11660* | 6.24 | 4.46 | 1.78 | phosphate-induced protein 1 conserved region domain containing protein, expressed |
|  | *LOC_Os06g11740* | 6.8 | 5.1 | 1.7 | OsAPRL2 adenosine 5'-phosphosulfate reductase-like OsAPRL2, expressed |
|  | *LOC_Os06g11860* | 4.93 | 3.5 | 1.43 | ethylene-responsive transcription factor, putative, expressed |
|  | *LOC_Os06g11990* | 5.34 | 6.77 | 1.43 | expressed protein |
|  | *LOC_Os06g12040* | 3.35 | 1.96 | 1.39 | mTERF family protein, expressed |
|  | *LOC_Os06g12170* | 3.61 | 0.06 | 3.55 | expressed protein |
|  | *LOC_Os06g12210* | 0 | 1.2 | 1.2 | helix-loop-helix DNA-binding domain containing protein, expressed |
| *PQ9* | *LOC_Os06g46340* | 0.7 | 3.13 | 2.43 | glycosyl hydrolase, family 31, putative, expressed |
|  | *LOC_Os06g46460* | 1.85 | 0 | 1.85 | expressed protein |
|  | *LOC_Os06g46500* | 0.15 | 2.25 | 2.1 | monocopper oxidase, putative, expressed |
|  | *LOC_Os06g46570* | 1.16 | 2.44 | 1.28 | galactosyltransferase, putative, expressed |
|  | *LOC_Os06g46680* | 0.59 | 2.03 | 1.44 | cytochrome P450, putative, expressed |
|  | *LOC_Os06g46754* | 4.11 | 5.22 | 1.11 | RIC10, putative, expressed |
|  | *LOC_Os06g46799* | 4.62 | 6.8 | 2.18 | peroxidase precursor, putative, expressed |
|  | *LOC_Os06g46930* | 6.34 | 4.64 | 1.7 | ribosomal protein L24, putative, expressed |
|  | *LOC_Os06g47000* | 3.06 | 4.47 | 1.41 | external NADH-ubiquinone oxidoreductase 1, mitochondrial precursor, putative, expressed |
|  | *LOC_Os06g47200* | 5.48 | 6.55 | 1.07 | LTPL85 - Protease inhibitor/seed storage/LTP family protein precursor, expressed |
|  | *LOC_Os06g47544* | 4.85 | 3.5 | 1.35 | expressed protein |
|  | *LOC_Os06g47600* | 0.72 | 1.88 | 1.16 | thaumatin family domain containing protein, expressed |
|  | *LOC_Os06g47760* | 3.24 | 1.82 | 1.42 | phytosulfokine receptor precursor, putative, expressed |
|  | *LOC_Os06g47820* | 0.15 | 1.53 | 1.38 | protein kinase domain containing protein, expressed |
|  | *LOC_Os06g47850* | 0.11 | 1.67 | 1.56 | zinc finger family protein, putative, expressed |
|  | *LOC_Os06g47970* | 5.09 | 3.93 | 1.16 | expressed protein |
| *PQ13* | *LOC_Os11g19210* | 0.03 | 1.31 | 1.28 | beta-D-xylosidase, putative, expressed |
|  | *LOC_Os11g19764* | 3.39 | 5 | 1.61 | expressed protein |
|  | *LOC_Os11g19800* | 1.97 | 0.68 | 1.29 | Ras family domain containing protein, expressed |
|  | *LOC_Os11g20170* | 2.78 | 4.37 | 1.59 | expressed protein |
|  | *LOC_Os11g20239* | 3.38 | 0.03 | 3.35 | expressed protein |
|  | *LOC_Os11g20310* | 3.91 | 0.2 | 3.71 | expressed protein |
|  | *LOC_Os11g20330* | 3.92 | 0 | 3.92 | expressed protein |
|  | *LOC_Os11g20384* | 3.25 | 0.92 | 2.33 | SacI homology domain containing protein, expressed |
|  | *LOC_Os11g22250* | 0 | 3.53 | 3.53 | expressed protein |
|  | *LOC_Os11g22350* | 2.94 | 0.6 | 2.34 | white-brown complex homolog protein, putative, expressed |
|  | *LOC_Os11g23790* | 5.69 | 2.7 | 2.99 | expressed protein |
|  | *LOC_Os11g24060* | 0.12 | 1.36 | 1.24 | permease domain containing protein, putative, expressed |
|  | *LOC_Os11g24070* | 0.36 | 1.92 | 1.56 | LTPL10 - Protease inhibitor/seed storage/LTP family protein precursor, expressed |
|  | *LOC_Os11g24140* | 7.28 | 2.65 | 4.63 | plastocyanin-like domain containing protein, putative, expressed |
|  | *LOC_Os11g24180* | 0.45 | 1.79 | 1.34 | OsSCP50 - Putative Serine Carboxypeptidase homologue, expressed |
|  | *LOC_Os11g24374* | 0.49 | 1.65 | 1.16 | OsSCP55 - Putative Serine Carboxypeptidase homologue, expressed |
|  | *LOC_Os11g24450* | 2.66 | 1.01 | 1.65 | mitochondrial carrier protein, putative, expressed |
|  | *LOC_Os11g24800* | 1.96 | 0.07 | 1.89 | expressed protein |
|  | *LOC_Os11g25080* | 3.81 | 0 | 3.81 | expressed protein |
|  | *LOC_Os11g25220* | 0 | 2.23 | 2.23 | oxidoreductase, short chain dehydrogenase/reductase domain containing protein, expressed |
|  | *LOC_Os11g25230* | 0 | 3.53 | 3.53 | tropinone reductase, putative |
|  | *LOC_Os11g25330* | 0 | 5.38 | 5.38 | nucleoside-triphosphatase, putative, expressed |
|  | *LOC_Os11g25454* | 2.56 | 1.27 | 1.29 | cytokinin-N-glucosyltransferase 1, putative, expressed |
|  | *LOC_Os11g25470* | 3.23 | 0 | 3.23 | expressed protein |
|  | *LOC_Os11g25720* | 0.81 | 2.14 | 1.33 | cytokinin-N-glucosyltransferase 1, putative, expressed |
|  | *LOC_Os11g26150* | 4.89 | 2.95 | 1.94 | expressed protein |
|  | *LOC_Os11g26340* | 2.31 | 3.66 | 1.35 | RALFL24 - Rapid ALkalinization Factor RALF family protein precursor, expressed |
|  | *LOC_Os11g26350* | 2.36 | 0 | 2.36 | expressed protein |
|  | *LOC_Os11g26570* | 0.21 | 1.54 | 1.33 | dehydrin, putative, expressed |
|  | *LOC_Os11g26720* | 0.3 | 2.33 | 2.03 | expressed protein |
|  | *LOC_Os11g26860* | 5.95 | 7.61 | 1.66 | serine hydroxymethyltransferase, mitochondrial precursor, putative, expressed |
|  | *LOC_Os11g26880* | 0.25 | 2.55 | 2.3 | RALFL14 - Rapid ALkalinization Factor RALF family protein precursor, expressed |
|  | *LOC_Os11g26990* | 0.25 | 3.2 | 2.95 | expressed protein |
|  | *LOC_Os11g27130* | 1.58 | 0 | 1.58 | HAT dimerisation protein, putative, expressed |
|  | *LOC_Os11g27240* | 1.64 | 0 | 1.64 | pentatricopeptide repeat domain containing protein, putative, expressed |
|  | *LOC_Os11g27400* | 0 | 2.09 | 2.09 | glycosyl hydrolase, putative, expressed |
|  | *LOC_Os11g27540* | 1.98 | 3.17 | 1.19 | CRP7 - Cysteine-rich family protein precursor, expressed |
|  | *LOC_Os11g27730* | 5.72 | 0 | 5.72 | cytochrome P450, putative, expressed |
|  | *LOC_Os11g27795* | 0.15 | 2.04 | 1.89 | expressed protein |
|  | *LOC_Os11g28104* | 0.12 | 6.9 | 6.78 | TKL_IRAK_DUF26-lf.4 - DUF26 kinases have homology to DUF26 containing loci, expressed |
|  | *LOC_Os11g28130* | 4.27 | 0 | 4.27 | expressed protein |
|  | *LOC_Os11g28310* | 0 | 1.03 | 1.03 | P21-Rho-binding domain containing protein, putative, expressed |
|  | *LOC_Os11g28430* | 3.39 | 0 | 3.39 | expressed protein |
| *PQ14* | *LOC_Os12g02300* | 0 | 4.68 | 4.68 | LTPL26 - Protease inhibitor/seed storage/LTP family protein precursor, expressed |
|  | *LOC_Os12g02310* | 1.52 | 5.15 | 3.63 | LTPL11 - Protease inhibitor/seed storage/LTP family protein precursor, expressed |

**Table S5** The haplotypes of 201 varieties at *LOC_Os06g11130*.

| POS | 5835826 | 5836263 | 5836330 | 5836463 | 5836905 | 5836978 | Variety name from 3k genome |
| --- | --- | --- | --- | --- | --- | --- | --- |
| Hap1 | C | A | C | C | C | T | CX250 |
| Hap1 | C | A | C | C | C | T | CX142 |
| Hap1 | C | A | C | C | C | T | B169 |
| Hap1 | C | A | C | C | C | T | B115 |
| Hap1 | C | A | C | C | C | T | CX139 |
| Hap1 | C | A | C | C | C | T | CX26 |
| Hap1 | C | A | C | C | C | T | B016 |
| Hap1 | C | A | C | C | C | T | CX75 |
| Hap1 | C | A | C | C | C | T | B162 |
| Hap1 | C | A | C | C | C | T | CX2 |
| Hap1 | C | A | C | C | C | T | B176 |
| Hap1 | C | A | C | C | C | T | B027 |
| Hap1 | C | A | C | C | C | T | CX286 |
| Hap1 | C | A | C | C | C | T | CX6 |
| Hap1 | C | A | C | C | C | T | B249 |
| Hap1 | C | A | C | C | C | T | B212 |
| Hap1 | C | A | C | C | C | T | CX150 |
| Hap1 | C | A | C | C | C | T | B011 |
| Hap1 | C | A | C | C | C | T | CX146 |
| Hap1 | C | A | C | C | C | T | CX79 |
| Hap1 | C | A | C | C | C | T | CX288 |
| Hap1 | C | A | C | C | C | T | B127 |
| Hap1 | C | A | C | C | C | T | CX99 |
| Hap1 | C | A | C | C | C | T | B079 |
| Hap1 | C | A | C | C | C | T | CX10 |
| Hap1 | C | A | C | C | C | T | CX207 |
| Hap1 | C | A | C | C | C | T | CX111 |
| Hap1 | C | A | C | C | C | T | CX91 |
| Hap1 | C | A | C | C | C | T | B225 |
| Hap1 | C | A | C | C | C | T | B165 |
| Hap1 | C | A | C | C | C | T | B219 |
| Hap1 | C | A | C | C | C | T | CX101 |
| Hap1 | C | A | C | C | C | T | CX318 |
| Hap1 | C | A | C | C | C | T | B129 |
| Hap1 | C | A | C | C | C | T | B215 |
| Hap1 | C | A | C | C | C | T | CX341 |
| Hap1 | C | A | C | C | C | T | B110 |
| Hap1 | C | A | C | C | C | T | CX88 |
| Hap1 | C | A | C | C | C | T | CX269 |
| Hap1 | C | A | C | C | C | T | CX274 |
| Hap1 | C | A | C | C | C | T | CX231 |
| Hap1 | C | A | C | C | C | T | CX233 |
| Hap1 | C | A | C | C | C | T | CX403 |
| Hap1 | C | A | C | C | C | T | CX230 |
| Hap1 | C | A | C | C | C | T | CX205 |
| Hap1 | C | A | C | C | C | T | CX225 |
| Hap1 | C | A | C | C | C | T | CX276 |
| Hap1 | C | A | C | C | C | T | CX64 |
| Hap1 | C | A | C | C | C | T | CX247 |
| Hap1 | C | A | C | C | C | T | B104 |
| Hap1 | C | A | C | C | C | T | B253 |
| Hap1 | C | A | C | C | C | T | CX329 |
| Hap1 | C | A | C | C | C | T | B254 |
| Hap1 | C | A | C | C | C | T | B151 |
| Hap1 | C | A | C | C | C | T | B096 |
| Hap1 | C | A | C | C | C | T | CX74 |
| Hap1 | C | A | C | C | C | T | B259 |
| Hap1 | C | A | C | C | C | T | B109 |
| Hap1 | C | A | C | C | C | T | B154 |
| Hap1 | C | A | C | C | C | T | B171 |
| Hap1 | C | A | C | C | C | T | B112 |
| Hap1 | C | A | C | C | C | T | B121 |
| Hap1 | C | A | C | C | C | T | CX155 |
| Hap1 | C | A | C | C | C | T | B102 |
| Hap1 | C | A | C | C | C | T | B229 |
| Hap1 | C | A | C | C | C | T | CX145 |
| Hap1 | C | A | C | C | C | T | CX306 |
| Hap1 | C | A | C | C | C | T | CX265 |
| Hap1 | C | A | C | C | C | T | CX317 |
| Hap1 | C | A | C | C | C | T | CX316 |
| Hap1 | C | A | C | C | C | T | CX70 |
| Hap1 | C | A | C | C | C | T | B086 |
| Hap1 | C | A | C | C | C | T | B268 |
| Hap1 | C | A | C | C | C | T | B210 |
| Hap1 | C | A | C | C | C | T | CX140 |
| Hap1 | C | A | C | C | C | T | CX235 |
| Hap1 | C | A | C | C | C | T | CX37 |
| Hap1 | C | A | C | C | C | T | B026 |
| Hap1 | C | A | C | C | C | T | CX157 |
| Hap1 | C | A | C | C | C | T | B167 |
| Hap1 | C | A | C | C | C | T | B089 |
| Hap1 | C | A | C | C | C | T | B093 |
| Hap1 | C | A | C | C | C | T | B032 |
| Hap1 | C | A | C | C | C | T | CX103 |
| Hap1 | C | A | C | C | C | T | CX346 |
| Hap1 | C | A | C | C | C | T | CX158 |
| Hap1 | C | A | C | C | C | T | CX148 |
| Hap1 | C | A | C | C | C | T | B173 |
| Hap1 | C | A | C | C | C | T | CX120 |
| Hap1 | C | A | C | C | C | T | CX534 |
| Hap1 | C | A | C | C | C | T | B263 |
| Hap1 | C | A | C | C | C | T | B252 |
| Hap1 | C | A | C | C | C | T | B233 |
| Hap1 | C | A | C | C | C | T | B024 |
| Hap1 | C | A | C | C | C | T | B258 |
| Hap1 | C | A | C | C | C | T | CX251 |
| Hap1 | C | A | C | C | C | T | CX356 |
| Hap1 | C | A | C | C | C | T | B006 |
| Hap1 | C | A | C | C | C | T | CX330 |
| Hap1 | C | A | C | C | C | T | CX390 |
| Hap1 | C | A | C | C | C | T | CX56 |
| Hap1 | C | A | C | C | C | T | CX359 |
| Hap1 | C | A | C | C | C | T | CX131 |
| Hap1 | C | A | C | C | C | T | B003 |
| Hap1 | C | A | C | C | C | T | B126 |
| Hap1 | C | A | C | C | C | T | B239 |
| Hap1 | C | A | C | C | C | T | CX133 |
| Hap1 | C | A | C | C | C | T | CX19 |
| Hap1 | C | A | C | C | C | T | CX315 |
| Hap1 | C | A | C | C | C | T | B090 |
| Hap2 | T | G | C | C | T | T | CX314 |
| Hap2 | T | G | C | C | T | T | B157 |
| Hap2 | T | G | C | C | T | T | CX96 |
| Hap2 | T | G | C | C | T | T | CX237 |
| Hap2 | T | G | C | C | T | T | CX352 |
| Hap2 | T | G | C | C | T | T | CX21 |
| Hap2 | T | G | C | C | T | T | CX97 |
| Hap2 | T | G | C | C | T | T | B033 |
| Hap2 | T | G | C | C | T | T | CX238 |
| Hap2 | T | G | C | C | T | T | CX85 |
| Hap2 | T | G | C | C | T | T | CX125 |
| Hap2 | T | G | C | C | T | T | CX8 |
| Hap2 | T | G | C | C | T | T | CX90 |
| Hap2 | T | G | C | C | T | T | B107 |
| Hap2 | T | G | C | C | T | T | B255 |
| Hap2 | T | G | C | C | T | T | CX219 |
| Hap2 | T | G | C | C | T | T | CX121 |
| Hap2 | T | G | C | C | T | T | B217 |
| Hap2 | T | G | C | C | T | T | CX144 |
| Hap2 | T | G | C | C | T | T | CX206 |
| Hap2 | T | G | C | C | T | T | CX73 |
| Hap2 | T | G | C | C | T | T | CX43 |
| Hap2 | T | G | C | C | T | T | CX278 |
| Hap2 | T | G | C | C | T | T | CX281 |
| Hap2 | T | G | C | C | T | T | CX296 |
| Hap2 | T | G | C | C | T | T | CX126 |
| Hap2 | T | G | C | C | T | T | CX134 |
| Hap2 | T | G | C | C | T | T | CX349 |
| Hap2 | T | G | C | C | T | T | CX154 |
| Hap2 | T | G | C | C | T | T | CX388 |
| Hap2 | T | G | C | C | T | T | CX303 |
| Hap2 | T | G | C | C | T | T | CX156 |
| Hap2 | T | G | C | C | T | T | CX376 |
| Hap2 | T | G | C | C | T | T | CX80 |
| Hap2 | T | G | C | C | T | T | CX347 |
| Hap2 | T | G | C | C | T | T | CX392 |
| Hap2 | T | G | C | C | T | T | CX33 |
| Hap2 | T | G | C | C | T | T | CX102 |
| Hap2 | T | G | C | C | T | T | CX93 |
| Hap2 | T | G | C | C | T | T | CX107 |
| Hap2 | T | G | C | C | T | T | CX92 |
| Hap2 | T | G | C | C | T | T | B106 |
| Hap2 | T | G | C | C | T | T | CX82 |
| Hap2 | T | G | C | C | T | T | CX84 |
| Hap2 | T | G | C | C | T | T | B214 |
| Hap2 | T | G | C | C | T | T | CX362 |
| Hap2 | T | G | C | C | T | T | CX363 |
| Hap3 | C | A | C | T | C | T | B008 |
| Hap3 | C | A | C | T | C | T | CX58 |
| Hap3 | C | A | C | T | C | T | CX396 |
| Hap3 | C | A | C | T | C | T | B170 |
| Hap3 | C | A | C | T | C | T | CX307 |
| Hap3 | C | A | C | T | C | T | CX389 |
| Hap3 | C | A | C | T | C | T | CX211 |
| Hap3 | C | A | C | T | C | T | CX397 |
| Hap3 | C | A | C | T | C | T | B004 |
| Hap3 | C | A | C | T | C | T | B160 |
| Hap3 | C | A | C | T | C | T | B005 |
| Hap3 | C | A | C | T | C | T | CX116 |
| Hap3 | C | A | C | T | C | T | CX350 |
| Hap3 | C | A | C | T | C | T | CX165 |
| Hap3 | C | A | C | T | C | T | CX391 |
| Hap3 | C | A | C | T | C | T | B269 |
| Hap3 | C | A | C | T | C | T | B240 |
| Hap3 | C | A | C | T | C | T | B235 |
| Hap3 | C | A | C | T | C | T | B168 |
| Hap4 | C | G | C | C | T | T | CX151 |
| Hap4 | C | G | C | C | T | T | CX3 |
| Hap4 | C | G | C | C | T | T | CX214 |
| Hap4 | C | G | C | C | T | T | B019 |
| Hap4 | C | G | C | C | T | T | CX266 |
| Hap4 | C | G | C | C | T | T | B025 |
| Hap4 | C | G | C | C | T | T | CX241 |
| Hap4 | C | G | C | C | T | T | CX243 |
| Hap4 | C | G | C | C | T | T | B246 |
| Hap4 | C | G | C | C | T | T | CX89 |
| Hap4 | C | G | C | C | T | T | CX32 |
| Hap4 | C | G | C | C | T | T | B095 |
| Hap4 | C | G | C | C | T | T | CX68 |
| Hap4 | C | G | C | C | T | T | CX141 |
| Hap4 | C | G | C | C | T | T | CX210 |
| Hap4 | C | G | C | C | T | T | CX270 |
| Hap4 | C | G | C | C | T | T | CX343 |
| Hap4 | C | G | C | C | T | T | CX83 |
| Hap4 | C | G | C | C | T | T | B091 |
| Hap5 | C | G | C | C | T | G | CX72 |
| Hap5 | C | G | C | C | T | G | CX143 |
| Hap5 | C | G | C | C | T | G | CX66 |
| Hap5 | C | G | C | C | T | G | CX355 |
| Hap6 | C | G | A | C | T | T | CX98 |
| Hap7 | T | G | C |  | T | T | CX122 |
